# A meta-review of DNA-based identification methods and mislabeling analysis of Eastern South Pacific seafood

**DOI:** 10.1101/2025.08.11.669513

**Authors:** Alan Marín

## Abstract

The Eastern South Pacific Ocean is a nutrient-rich and highly diverse region that plays a pivotal role in the global seafood trade. Despite its importance, the seafood industry in this region is hindered by complex supply chains and insufficient regulation frameworks, which facilitate illegal practices, including mislabeling. DNA-based methods have emerged as essential tools for seafood authentication, helping to mitigate mislabeling and supporting conservation strategies. This study provides the first comprehensive review of DNA-based methods used globally to identify seafood species from the Eastern South Pacific Ocean. Historical and current techniques were systematically examined, with a focus on emerging technologies that offer promising applications in the field. Nearly three decades of research have enabled the successful identification of over 200 commercially valuable species using more than ten distinct DNA-based methods. Fish and mollusks constituted the most extensively studied seafood groups, with DNA sequencing emerging as the predominant technique. Furthermore, a global mislabeling meta-analysis, encompassing 1,806 seafood products from the Eastern South Pacific Ocean, revealed an estimated mislabeling rate of 24.8% (95% CI [22.9-26.9]). Notably, a substantial proportion of mislabeled and substituted products corresponds to highly threatened shark species. This raises serious conservation concerns, particularly given that nations bordering the Eastern South Pacific Ocean are major players in the global shark trade. Overall, the findings of this review underscore the urgent need to integrate advanced DNA-based techniques into existing regulatory frameworks. They also establish a solid foundation for developing targeted policies and encouraging collaborative efforts among nations in this region.

## 1. Introduction

Fish and seafood products have become some of the most traded commodities around the world, with an estimated annual production value of US$ 472 billion (Hellberg, 2024; Lawrence et al., 2022; FAO, 2024). During 2022, seafood contributed an estimated 20.7 kg per capita and provided about 15% of the world’s animal protein supply (FAO, 2024). A significant portion of this production comes from the coastal and oceanic fisheries of the eastern South Pacific Ocean (hereafter referred to as ESPO). In 2021, landings from the ESPO reached 10 million tons, accounting for 12.5% of global landings (FAO, 2024). Marine ecosystems in the ESPO are characterized by nutrient-rich waters and highly productive coastal upwelling zones. These factors make the ESPO one of the most productive marine systems in the world, exhibiting extremely high primary productivity that supports a rich and diverse array of marine life (Gonzalez et al., 2019; Serratosa et al., 2020).

Despite management efforts, several fisheries in developing countries, including those bordering the ESPO region, suffer from weak regulatory systems, wildlife trafficking, and high levels of illegal, unreported, and unregulated (IUU) fishing (Marín et al., 2018; De la Puente et al., 2020; Gozzer-Wuest et al., 2023). These issues create opportunities for deceptive practices such as mislabeling and species substitution throughout the supply chain (Lawrence et al., 2022). Seafood mislabeling not only undermines conservation and sustainability initiatives, but it also poses risks to consumers’ health, economic well-being, trust, and even religious beliefs (Marín et al., 2013; Silva & Hellberg, 2021; Neo et al., 2022).

In this context, accurate species identification of commercially important marine resources is a crucial step toward effective management and prevention of dishonest practices. Although some seafood species are easily recognized, many marine organisms undergo various adaptive evolutionary processes that complicate their morphological differentiation. These processes include convergent phenotypic evolution, interspecific conservatism, phenotypic plasticity, hybridization, and morphological variation within species (Pérez-Jiménez et al., 2005; Hanner et al., 2011; Mynhardt et al., 2014; Pliego-Cárdenas et al., 2016; Neves et al., 2021; Marín et al., 2022). Furthermore, most processed and cooked seafood products lack distinctive morphological features, making them unsuitable for identification through conventional taxonomic methods. To address this issue, several alternative molecular identification methods have been developed, which are independent of morphological characters and have proven to be effective for identifying both fresh and processed seafood species (Ali et al., 2012; Lago et al., 2014).

Recent advances in molecular biology have led to the development of various accurate and sensitive methods that can of identify organisms at the species level, even from highly processed samples or minuscule amounts of tissue. Consequently, a range of analytical techniques based on proteins and DNA have been employed to authenticate seafood products (Ali et al., 2012; Lago et al., 2014; Silva & Hellberg, 2021). However, the heat-labile nature of most proteins limits the applicability of protein-based techniques to fresh or lightly processed seafood samples (Teletchea, 2009; Silva & Hellberg, 2021; Aditi et al., 2024). In contrast, DNA is more resistant to heat degradation, making it suitable for analyzing highly processed seafood products, such as canned or cooked items. This advantage allows DNA techniques to overcome the limitations associated with protein-based methods (Teletchea, 2009). Furthermore, DNA can be PCR-amplified from very small amounts of starting material, which can be collected from virtually any organic substrate, including fins, blood, mucus, and bones (Teletchea, 2009; Silva & Hellberg, 2021; Aditi et al., 2024).

Although some comprehensive reviews have discussed DNA-based techniques for global seafood authentication and mislabeling (Teletchea, 2009; Ali et al., 2012; Lago et al., 2014; Pardo et al., 2016; Luque & Donlan, 2019; Silva & Hellberg, 2021; Aditi et al., 2024), most of the reviewed studies from South American countries are primarily focused on research conducted in Brazil, which examines species from the western South Atlantic Ocean. In contrast, reviewed studies addressing seafood species from the ESPO region remain underrepresented. There is a noticeable lack of reviews on DNA detection of seafood diversity and mislabeling rates in articles from ESPO nations, despite the regiońs rich marine biodiversity, significant wild-capture and mariculture production, and the wide variety of seafood products that these nations offer commercially around the globe.

This review provides a comprehensive overview of both past and present DNA-based methods used to identify commercially important species from the ESPO region. It offers a summary of the techniques, DNA markers, and oligonucleotide primers found in the reviewed literature. This review also intends to examine the ESPO’s species diversity that has been identified using DNA methods, presenting the most current database of commercially significant species accompanied by molecular identification records. Furthermore, I performed an analysis of their current conservation statuses based on information from the International Union for Conservation of Nature (IUCN). Lastly, a systematic meta-review of seafood mislabeling was conducted based on data sourced from the surveyed articles, which feature samples collected from both domestic and international markets where ESPO species are sold.

## 2. Materials and methods

### 2.1 Data collection and inclusion criteria

This study is based on an exhaustive analysis of peer-reviewed papers published between mid-December 1998 and early July 2025. The Level 1 screening focused on articles that report the development and application of DNA-based tools for identifying commercial marine samples from fisheries and mariculture within the ESPO region. Samples were collected throughout the seafood supply chain, including fish landings, farms, curing facilities, wholesale fish markets, manufacturers, fishmongers, local markets, supermarkets, ethnic markets, restaurants, and exporters in ESPO’s countries and international markets. Articles that originated outside the ESPO’s countries were included only if they analyzed products that contained declared species from the ESPO region, specifically those identified as imported commodities. Relevant research articles were sourced from multiple databases, including Google Scholar, Direct, PubMed, ResearchScience, SciELO, and Scopus, using the following search terms (in both English and Spanish): “canned” OR “crab” OR “crustaceans” OR “fish” OR “fish market” OR “gastropods” OR “DNA barcoding” OR “DNA meta-barcoding” OR “DNA-based techniques” OR “DNA-sequencing” OR “High-throughput sequencing” OR “mislabeling” OR “molecular identification” OR “molecular markers” OR “mollusks” OR “polymerase chain reaction (PCR)” OR “protected marine species” OR “qPCR” OR “seafood authentication” OR “seafood fraud” OR “seafood supply chain” OR “sharks” OR “shellfish” OR “species identification” OR “species-specific primers” OR “Eastern South Pacific” OR “South America” OR “Chile” OR “Colombia” OR “Ecuador” OR “Peru”. Cited references from the identified articles were also reviewed to find related papers not identified in the initial search.

The most targeted studies focused on developing molecular markers and applying DNA-based identification tools for seafood authentication and detecting mislabeling. However, studies that utilized DNA-based methods for identifying new range expansions, species occurrences, phylogenetic relationships, or population genetics were considered only if the specimens analyzed had commercial origins or were captured by fishermen, indicating the type of retail venue and location where the samples were obtained. Studies based on self-collected samples from wild populations and environmental DNA (eDNA) were excluded. However, articles validating novel DNA-based methods (e.g. assessing the performance of novel primers) using self-collected samples from natural populations were included if the target species had economic significance. Unpublished studies, preprints, abstracts, or any research that had not undergone a peer-review process were disregarded. Additionally, this review included studies that analyzed samples from protected or endangered species of commercial value in local or international markets (whether these samples were commercially obtained or seized by authorities) whose trade is banned or regulated. Lastly, this review also examined articles that reported on the molecular authentication of fish feed claimed to contain commercially valuable species from the ESPO.

### 2.2 DNA-based methods, molecular markers, and primers

For practical purposes, DNA-based methods used for seafood authentication of ESPO’s species were categorized into three distinct groups. The first category consists of methods that rely on the amplification of DNA by an enzymatic reaction. The results from these methods are based on the visualization of a specific size fragment or band pattern in a gel matrix, a positive colorimetric reaction, or a fluorescence signal pattern displayed digitally. This category includes the following methods: 1) PCR-RFLP (Restriction Fragment Length Polymorphism), 2) PCR-RAPD (Random Amplified Polymorphic DNA), 3) PCR-SSCP (Single Strand Conformation Polymorphism), 4) PCR-SSP (Species Specific Primers), 5) qPCR-SSP, 6) Multiplex PCR and qPCR, 7) qPCR High-Resolution Melting (HRM), and 8) PCR-Microarrays, and 9) Loop-Mediated Isothermal Amplification (LAMP). The second category encompasses methods that depend on DNA sequencing technologies. This category further divides into two subcategories: a) sequences generated through a PCR step followed by the Sanger Sequencing technique, which includes: 10) FINS (Forensically Informative Nucleotide Sequencing) and DNA Barcoding; and b) sequences generated by high-throughput NGS (Next Generation Sequencing) technologies, specifically 11) HTS NGS-metabarcoding. The third category includes seafood species identification based on 12) mass spectrometry analysis. The genetic markers discussed in the reviewed articles were organized into three levels based on 1) genome origin: mitochondrial, nuclear, and random markers (like those targeted by RAPD primers), 2) target organisms, and 3) DNA-based methods. Similarly, all oligonucleotide primers reported in the analyzed articles were classified into two levels according to 1) target molecular marker (e.g., mitochondrial COI gene, nuclear rhodopsin gene) and 2) targeted animal groups.

### 2.3 Commercial species diversity and conservation statuses

The species composition of commercially traded seafood from the ESPO region was assessed, considering both domestic and international markets. This assessment was based on the species identification results reported in the reviewed articles. This review focused exclusively on seafood species from the ESPO region, which includes marine species (such as sharks and tuna) and diadromous fish (like eel, herring, salmon, and trout). Strict freshwater species (such as tilapia) were not included in the analysis. Only species that are native to the ESPO region were considered; however, introduced or naturalized species of commercial value were also taken into account. Species identified in reviewed studies collected from domestic ESPO markets, which have a confirmed distribution range outside the ESPO (likely sourced from imported commodities), were excluded from the assessment. Distribution ranges were verified using several databases: The Ocean Biodiversity Information System OBIS (https://obis.org), FishBase (https://fishbase.org), SeaLifeBase (https://www.sealifebase.org), The Barcode of Life Data System -BOLD-(https://boldsystems.org), and the World Register of Marine Species -WoRMS- (https://marinespecies.org). All identified species reported in the analyzed studies were organized at the taxonomic resolution level based on the current accepted classification and scientific names provided by the WoRMS database. Additionally, the current conservation status of all species identified in the papers analyzed in this review were searched in the IUCN Red List of Threatened Species website (https://www.iucnredlist.org).

### 2.4 Mislabeling meta-analysis

The mislabeling meta-analysis dataset was constructed using articles that met the Level 1 screening criteria. These articles underwent a second screening based on Level 2 criteria, which required that the study analyzes samples of commercial origin. Following this, a third screening was conducted using the Level 3 criteria, which mandated that detailed information for each analyzed sample be provided either in the main text or as a supplementary file. This information included the label (commercial name), type of product (e.g. fresh, cooked, cured, canned), and sampling venue (e.g. fish landing site, market). Only samples identified at the species level were considered, and classifications above the species level were not included. The samples included in the mislabeling meta-analysis consisted of seafood products that were labeled as containing an ESPO species, such as “Peruvian anchoveta” (*Engraulis ringens*). These products were either identified as the correct species or were found to be substituted with another ESPO species (e.g. *Sardinops sagax*), a non-ESPO species (e.g. “European anchovy”, *E. encrasicolus*), or a freshwater species. Additionally, the analysis included cases where ESPO species were used to replace non-native ESPO species from international markets, for example, the Peruvian scallop (*Argopecten purpuratus*) being sold as the European king scallop (*Pecten maximus*).

In an attempt to recover samples that had been identified at a higher taxonomic level due to a lack of reference sequences at the time of publication, a curation update step was implemented. This involved using publicly available DNA sequences cited in the reviewed papers, which were re-analyzed and compared against current reference sequences from the GenBank and BOLD databases.

To prevent the inclusion of samples that might have been classified as mislabeled due to a strict interpretation of expected common names, I carefully checked the label information (commercial designation or common name) and the reported scientific name of each analyzed sample. This information was compared against common names used in official landing reports, governmental websites, relevant literature, the FAO ASFIS database, and listed common names from the FishBase database. For instance, samples from domestic ESPO markets that were originally reported as mislabeled because they used generic names such as squid (“*calamar*”), flounder (“*lenguado*”), grouper (“*mero*”), shark (“*tiburón*” or “*tollo*”) were not considered mislabeled in this review if the identified species belonged to a seafood group implied by the declared generic term. This criterion was adopted because generic or umbrella market names are commonly used in seafood markets in the ESPO region, where no official lists of commercial designations for seafood species have been established. In fact, several acceptable market names included in the FDÁs The Seafood List (available at https://www.hfpappexternal.fda.gov/scripts/fdcc/index.cfm?set=SeafoodList) are examples of generic seafood names (e.g. abalone, anchovy, grouper, sea bass, shark). In contrast, for samples purchased within European markets, which have stricter seafood labeling regulations, products must be labeled with unique commercial designations and the scientific name, according to Regulation (EU) N° 1379/2013. Therefore, official market names were verified using the commercial and scientific name of seafood species listed on the European Union (EU) website (https://oceans-and-fisheries.ec.europa.eu/fisheries/markets-and-trade/seafood-markets/commercial-and-scientific-name-species_en). For samples collected in Canada, the Canadian Food Inspection Agency (CFIA) fish list was consulted from its official website (https://inspection.canada.ca/en/food-labels/labelling/industry/fish/list).

Cured samples from selected articles were pooled across studies and organized into different levels including label information, seafood type (e.g. finfish, sharks, bivalves), sampling venue, and country. The mislabeling rate was calculated using the formula [mn/n*100], where “mn” is the number of mislabeled samples and “n” is the total number of samples.

#### 2.4.1 Statistical analyses

Confidence intervals of the mislabeling rates were calculated using the “binom.confint” function from the “binom” package implemented in R (R for Mac OS X GUI version 4.2.3) (R core Team, 2022) to generate upper and lower estimates of the predicted mislabeling rates. The Agresti-Coull method was used for calculating 95% confidence intervals (CI) for large sample sizes (n > 40), while the Wilson method was chosen for small sample sizes (n ≤ 40) (Brown et al., 2001). Since these values represent percentages, any negative confidence intervals were reported as zero, and those exceeding 100% were reported as 100% (Bubernick & Faulstick, 1993). Exploratory analyses were conducted to assess the independence of the association between mislabeling rates based on continent, country, distribution channel, and seafood type. Categories with sample sizes of fewer than 25 (n < 25) were excluded from the analyses. The analyses utilized the Chi-square test or Fisher’s exact test, depending on whether any cell count was below 5. This approach was taken because Chi-square test relies on approximations that become unreliable with small expected cell counts (Kim, 2017). When a significant association between variables was observed (*p* < 0.05), further pairwise tests were conducted in R to determine which pairwise comparisons were significantly different. *p* values were adjusted for multiple comparisons using the Bonferroni correction method, and the α-level was set at 0.05 for all analyses. All R scripts used in the analyses are provided in Supplementary File S1 to S5.

## 3. Results and discussion

### 3.1 Data collection overview

This review compiled all the DNA-based methods used globally to identify commercially important species in the ESPO region. A total of 75 published articles (see Table 1) met the Level 1 screening criteria outlined in section 2.1 and were selected for analysis. The 75 reviewed articles were published across 43 different peer-review journals, with publication dates ranging from mid-December 1998 to early July 2025. Notably, there were no relevant articles published between 1999 and 2001. The journals “*Food Control*” and “*Journal of Agricultural and Food Chemistry*” were the most frequent publication venues, with 12 articles (16%) and 6 articles (8%), respectively. English was the predominant language, appearing in 69 articles (92%). Three articles were written in Spanish, and one article each was published in Chinese, German, and Korean. Out of the 75 studies, 30 were conducted entirely in ESPO countries: Chile (14 studies), Peru (12 studies), Colombia (3 studies), and Ecuador (1 study). The remaining 45 studies analyzed seafood samples collected around the world, including those from the ESPO region. These were conducted in Spain (17 studies), Germany (4), Japan (4), Mexico (4), Canada (3), China (2), France (2), Italy (2), and one study was carried out in each of the following countries: Austria, Brazil, Hong Kong, South Korea, Poland, Switzerland, and the United Kingdom. A total of 63 articles reported the authentication of commercial products from the ESPO. The remaining 12 articles validated their molecular assays using reference samples of non-commercial origin, which were obtained from natural populations within the ESPO, collected by researchers, donated by colleagues, or obtained from research facilities. Out of the 63 articles that reported the use of commercial ESPO products, 24 did not meet the Level 3 screening criteria, as they did not provide detailed information about each individual sample. This lack of detail prevented me from using their results in further mislabeling meta-analysis. As a result, only the findings from 39 papers were included in the mislabeling meta-analysis review (section 3.7).

**Table 1.** List of articles reviewed in this study.

| N° | Authors | Publication year | Journal (Language) | Target species | Method | DNA markers | ESPO seafood species type (supply channel and country of origin) | Conducted in | Passed screening Levels | Samples (n) in meta-analysis | Reference |
| --- | --- | --- | --- | --- | --- | --- | --- | --- | --- | --- | --- |
| 1 | Toro | 1998 | Journal of Shellfish Research (English) | Mussels | PCR-RFLP | ITS, PAMP | Wild (Chile) | CANADA | 1 | — | Toro (1998) |
| 2 | Rego et al. | 2002 | Journal of Agricultural and Food Chemistry (English) | Mussels | PCR-RAPD | Random | Wild (Chile) | SPAIN | 1 | — | Rego et al. (2002) |
| 3 | Rehbein | 2002 | Journal of Aquatic Food Product Technology (English) | Fish | PCR-RFLP SSCP | CYTB | Commercial (IFOMA Germany, imported from Chile and Peru) | GERMANY | 1 and 2 | — | Rehbein (2002) |
| 4 | Jérôme et al. | 2003a | Journal of Agricultural and Food Chemistry (English) | Sardines | PCR-RFLP | CYTB | Commercial (supermarket in France, canned in Peru) | FRANCE | 1, 2, and 3 | 1 | Jérôme et al. (2003a) |
| 5 | Jérôme et al. | 2003b | Journal of Agricultural and Food Chemistry (English) | Sardines | FINS | CYTB | Commercial (supermarket in France, origin Chile and Peru) | FRANCE | 1, 2, and 3 | 6 | Jérôme et al. (2003b) |
| 6 | Perez and García-Vazquez | 2004 | Journal of Food Protection (English) | Hake | PCR-RFLP | 5S rDNA, CYTB | Wild (Chile) | SPAIN | 1 | — | Perez and García-Vazquez (2004) |
| 7 | Toro et al. | 2005 | Journal of Shellfish Research (English) | Mussels | PCR-RFLP | ITS, PAMP | Wild (Chile) | CHILE | 1 | — | Toro et al. (2005) |
| 8 | Olivares-Paz et al. | 2006 | Investigaciones Marinas (Spanish) | Keyhole limpets | PCR-RFLP | CYTB | Wild (Chile) | SPAIN/CHILE | 1 | — | Olivares-Paz et al. (2006) |
| 9 | Marín et al. | 2007 | Journal of Shellfish Research (English) | Abalone | PCR-RAPD | RANDOM | Research facility (Chile) | CHILE | 1 | — | Marín et al. (2007) |
| 10 | Aguilera-Muñoz et al. | 2008 | Gayana (English) | Mollusks | PCR and band size polymorphism | ITS1, ITS2 | Commercial (Chile, Mexico) | CHILE | 1 and 2 | — | Aguilera-Muñoz et al. (2008) |
| 11 | Hernández et al. | 2008 | Journal of Fish Biology (English) | Shark fins | Multiplex PCR-SSP | ITS2 | Commercial (drying facility, commercial storage warehouse in Chile) | CHILE | 1, 2, and 3 | 200 | Hernández et al. (2008) |
| 12 | Espiñeira et al. | 2009a | Journal of Agricultural and Food Chemistry (English) | Bivalves | FINS | 18S rDNA | Wild (Chile, Peru) | SPAIN | 1 | — | Espiñeira et al. (2009a) |
| 13 | Espiñeira et al. | 2009b | European Food Research and Technology (English) | Salmon, trout, bream | FINS, PCR-RFLP | CYTB | Wild (Peru) and commercial (supermarket, Chile) | SPAIN | 1 and 2 | — | Espiñeira et al. (2009b) |
| 14 | Hernández et al. | 2010 | Hidrobiológica (English) | Shark fins | PCR-SSP | ITS2 | Commercial (storage warehouse, Chile) | CHILE | 1 and 2 | — | Hernández et al. (2010) |
| 15 | Tzika et al. | 2010 | Conservation Genetics (English) | Cetaceans | DNA sequencing | CYTB | Commercial (markets, Peru) | SWITZERLAND/ PERU | 1 and 2 | — | Tzika et al. (2010) |
| 16 | Aranceta-Garza et al. | 2011 | Food Control (English) | Abalone | PCR-SSCP | 18S rDNA | Commercial and wild (Retail market in Mexico, imported from Chile and Peru) | MEXICO | 1, 2, and 3 | 2 | Aranceta-Garza et al. (2011) |
| 17 | Fernández-Tajes et al. | 2011 | European Food Research and Technology (English) | Mussels | PCR-RFLP | PAPM, COI | Commercial and wild (Spanish supermarket, imported from Chile) | SPAIN | 1, 2, and 3 | 80 | Fernández-Tajes et al. (2011) |
| 18 | Ha et al. | 2011 | Journal of Agricultural and Food Chemistry (English) | Abalone | FINS | 16S rRNA, (COI was used as auxiliary marker) | Commercial (wholesale and retail outlets in Hong Kong, imported from Chile) | HONG KONG/UK | 1, 2, and 3 | 2 | Ha et al. (2011) |
| 19 | Herrero et al. | 2011 | Food Chemistry (English) | Salmon | qPCR-SSP | ITS1 | Commercial (supermarket in Chile) | SPAIN | 1 and 2 | — | Herrero et al. (2011) |
| 20 | Ardura et al. | 2012 | Fisheries Research (English) | Fish | DNA sequencing | 16S rRNA | Commercial (Spanish producers, species origin: Peru) | SPAIN | 1, 2, and 3 | 2 | Ardura et al. (2012) |
| 21 | Caballero et al. | 2012 | Molecular Ecology Resources (English) | Sharks | Multiplex PCR-SSP, DNA sequencing | COI, ITS2 | Commercial (landings in Colombia) | COLOMBIA | 1 and 2 | — | Caballero et al. (2012) |
| 22 | Haye et al. | 2012 | Food Control (English) | Crabs | DNA sequencing | COI | Commercial (seafood market and food store in Chile) | CHILE | 1 and 2 | — | Haye et al. (2012) |
| 23 | Näumann et al. | 2012 | Informationen aus der Fischereiforschung (German) | Scallops | DNA sequencing | 16S rRNA | Commercial (Imported or local commerce in Germany, imported from Peru) | GERMANY | 1, 2, and 3 | 1 | Näumann et al. (2012) |
| 24 | Horreo et al. | 2013 | Journal of the Science of Food and Agriculture (English) | Several animal species | DNA sequencing | 16S rRNA | Commercial (imported from Peru) | SPAIN | 1, 2, and 3 | 1 | Horreo et al. (2013) |
| 25 | Marín et al. | 2013 | Food Control (English) | Scallops | Multiplex PCR-SSP | 16S rRNA | Commercial (landings and market in Peru) | JAPAN | 1, 2, and 3 | 3 | Marín et al. (2013) |
| 26 | Chairi and Rebordinos | 2014 | Journal of Agricultural and Food Chemistry (English) | Anchovies | Multiplex PCR-SSP | 5S rDNA | Wild (Chile) | SPAIN | 1 | — | Chairi and Rebordinos (2014) |
| 27 | Santaclara et al. | 2014 | Food Chemistry (English) | Ling fish | FINS | COI | Wild (Chile) | SPAIN | 1 | — | Santaclara et al. (2014) |
| 28 | Marín et al. | 2015 | Fisheries Science (English) | Scallops | DNA sequencing | 16S rRNA | Commercial (landings in Peru) | JAPAN | 1, 2, and 3 | 8 | Marín et al. (2015) |
| 29 | Velez-Zuazo et al. | 2015 | Fisheries Research (English) | Sharks | DNA sequencing | COI | Commercial (landings in Peru) | PERU | 1 and 2 | — | Velez-Zuazo et al. (2015) |
| 30 | Galal-Khallaf et al. | 2016 | Food Chemistry (English) | Shrimps/ Cephalopods | DNA sequencing | COI, 16S rRNA | Commercial (supermarkets in Spain and Egypt, imported from Ecuador and Chile) | SPAIN/EGYPT | 1, 2, and 3 | 18 | Galal-Khallaf et al. (2016) |
| 31 | Pliego Cárdenas et al. | 2016 | American Malacological Bulletin (English) | Octopus | DNA sequencing | COI, COIII, 16S rRNA | Commercial (artisanal fishery in Ecuador) | MEXICO/ ECUADOR | 1, 2, and 3 | 7 | Pliego Cárdenas et al. (2016) |
| 32 | Sanchez et al. | 2016 | Fisheries Science (English) | Cephalopods | DNA sequencing | 16S rRNA | Wild (Peru) | JAPAN | 1 | — | Sanchez et al. (2016) |
| 33 | Velasco et al. | 2016 | Heliyon (English) | Anchovies | FINS | CYTB | Commercial (markets, supermarkets, and specialized stores in Spain, origin Chile and Peru) | SPAIN | 1, 2, and 3 | 11 | Velasco et al. (2016) |
| 34 | Abbadi et al. | 2017 | Journal of the Science of Food and Agriculture (English) | Bivalves | HTS Pyroseq | COI, 16S rRNA | Commercial (customs premises, market in Italy, imported from Chile) | ITALY | 1, 2, and 3 | 44 | Abbadi et al. (2017) |
| 35 | Jilberto et al. | 2017 | Food Chemistry (English) | Mussels | HRM, RFLP | PAPM | Wild (Chile) | SPAIN | 1 | — | Jilberto et al. (2017) |
| 36 | Marín et al. | 2017 | Aquaculture Research (English) | Scallops | Multiplex PCR-SSP | 16S rRNA | Commercial (landings in Peru and supermarket in UK, imported from Peru) | JAPAN | 1, 2, and 3 | 65 | Marín et al. (2017) |
| 37 | Canales-Aguirre et al. | 2018 | Revista de Biología Marina y Oceanografía (English) | Fish | DNA sequencing | COI | Commercial (artisanal fishery in Chile) | CHILE | 1 and 2 | — | Canales-Aguirre et al. (2018) |
| 38 | Marín et al. | 2018 | Plos One (English) | Crustacean, fish, gastropods, mollusks | DNA sequencing | COI, 12S rRNA, 16S rRNA, Control Region | Commercial (landings, wholesales, markets, supermarkets, restaurants in Peru) | PERU | 1, 2, and 3 | 120 | Marín et al. (2018) |
| 39 | Mora et al. | 2018 | Salud y Tecnología Veterinaria (Spanish) | Fish | qPCR-SSP | ATP6, Control Region | Commercial (supermarkets in Peru) | PERU | 1 and 2 | — | Mora et al. (2018) |
| 40 | Shehata et al. | 2018 | Food Control (English) | Fish | DNA sequencing | COI | Commercial (Canadian Food and Inspection Agency surveillance) across Canada, imported from Chile and Peru) | CANADA | 1, 2, and 3 | 9 | Shehata et al. (2018) |
| 41 | Giusti et al. | 2019 | Food Control (English) | Anchovies | DNA sequencing | CYTB | Commercial (large-scale retail distribution in Italy, imported from Chile) | CHILE | 1, 2, and 3 | 9 | Giusti et al. (2019) |
| 42 | Hu et al. | 2019 | Science and Technology of Food Industry (Chinese) | Sea cucumber | DNA sequencing | COI, 16S rRNA | Commercial (supermarket in China, species origin: Peru) | CHINA | 1, 2, and 3 | 1 | Hu et al. (2019) |
| 43 | Larraín et al. | 2019 | Scientific Reports (English) | Mussels | RFLP | COI, 16S rRNA, PAPM, ITS, MAC-1, 18S rDNA | Wild (Chile) | CHILE | 1 | — | Larraín et al. (2019) |
| 44 | Shehata et al. | 2019 | Food Research International (English) | Fish | DNA sequencing | COI | Commercial (Canadian Food and Inspection Agency surveillance across Canada, imported from Chile, Colombia, Ecuador and Peru) | CANADA | 1, 2, and 3 | 12 | Shehata et al. (2019) |
| 45 | Yu et al. | 2019 | Journal of Food Hygiene and Safety (Korean) | Cephalopods | DNA sequencing | COI, 16S rRNA | Commercial (market in South Korea, imported from Chile and Peru) | KOREA | 1, 2, and 3 | 5 | Yu et al. (2019) |
| 46 | Biffi et al. | 2020 | Scientific Reports (English) | Cephalopods , fish, cetaceans | DNA sequencing | COI | Commercial (landings, wholesale, retail, supermarket, restaurants in Peru) | PERU | 1, 2, and 3 | 413 | Biffi et al. (2020) |
| 47 | Cabanillas-Torpoco et al. | 2020 | Zootaxa (English) | Sawfish | DNA sequencing | COI | Commercial (landings in Peru) | PERU | 1 and 2 | — | Cabanillas-Torpoco et al. (2020) |
| 48 | Colihueque et al. | 2020 | Recent Patents on Food, Nutrition & Agriculture (English) | Mussels | PCR-RFLP | 18S rDNA | Commercial (supermarkets in Chile) | CHILE | 1, 2, and 3 | 95 | Colihueque et al. (2020) |
| 49 | Prida et al. | 2020 | Foods (English) | Salmon, trout | PCR-RFLP | CYTB | Commercial (restaurants in Chile) | CHILE | 1, 2, and 3 | 84 | Prida et al. (2020) |
| 50 | Cañedo-Apolaya et al. | 2021 | Diversity (English) | Angel shark | DNA sequencing | COI | Commercial (landings and wholesale in Peru) | PERU | 1 and 2 | — | Cañedo-Apolaya et al. (2021) |
| 51 | Costa et al. | 2021 | Molecular Phylogenetics and Evolution (English) | Squids | DNA sequencing | COI, 16S rRNA, Rhodopsin | Commercial (markets in Ecuador and Peru) | BRAZIL | 1 and 2 | — | Costa et al. (2021) |
| 52 | Gense et al. | 2021 | Foods (English) | Bivalves | NHT NGS-Metabar-coding | 16S rRNA | Commercial (delicacy shop in Austria, imported from Peru) | AUSTRIA | 1, 2, and 3 | 16 | Gense et al. (2021) |
| 53 | Kappel et al. | 2021 | Analytical and Bioanalytical Chemistry (English) | *Penaeus* shrimps | Microarray, DNA sequencing | COI, 16S rRNA | Commercial (retailer in Germany, declared origin: Ecuador) | GERMANY | 1, 2, and 3 | 1 | Kappel et al. (2021) |
| 54 | Klapper and Schröder | 2021 | Food Control (English) | Scallops | Multiplex qPCR, DNA sequencing | 16S rRNA | Commercial (artisanal fishery in Chile) | GERMANY | 1, 2, and 3 | 2 | Klapper and Schröder (2021) |
| 55 | Marín et al. | 2021 | Journal for Nature Conservation (English) | Fish | qPCR-SSP, DNA sequencing | COI, 16S rRNA | Commercial (landings, wholesale, and markets in Peru) | PERU | 1, 2, and 3 | 61 | Marín et al. (2021) |
| 56 | Parrondo et al. | 2021 | Food Control (English) | Scallops | DNA sequencing | 16S rRNA | Commercial (fish shops, supermarkets, and restaurants in Spain imported from Chile or Peru) | SPAIN | 1, 2, and 3 | 77 | Parrondo et al. (2021) |
| 57 | Quintrel et al. | 2021 | Foods (English) | Mussels | PCR-RFLP, qPCR-HRM | PAPM | Wild and commercial (Chile) | CHILE | 1 and 2 | — | Quintrel et al. (2021) |
| 58 | Velez-Zuazo et al. | 2021 | Food Control (English) | Fish | DNA sequencing | COI | Commercial (fish markets, restaurants, and supermarkets in Peru) | PERU | 1 and 2 | — | Velez-Zuazo et al. (2021) |
| 59 | Xing et al. | 2021 | Food Control (English) | Sea cucumber | DNA sequencing | COI, 16S rRNA | Commercial (supermarket in China, origin: Peru) | CHINA | 1, 2, and 3 | 1 | Xing et al. (2021) |
| 60 | Asorey et al. | 2022 | Food Chemistry: Molecular Sciences (English) | Mussels | HRM, FINS | PAPM, H1C | Commercial (growing center in Chile) | CHILE | 1, 2, and 3 | 61 | Asorey et al. (2022) |
| 61 | Dufflocq et al. | 2022 | Food Control (English) | Swordfish | DNA sequencing | COI | Commercial (fish market, open-air market, and Supermarkets in Chile) | CHILE | 1, 2, and 3 | 47 | Dufflocq et al. (2022) |
| 62 | Giusti et al. | 2022 | Food Control (English) | Mussels | PCR-RFLP | PAPM | Commercial (national large retailers, ethnic shop (Italy, imported from Chile) | ITALY | 1, 2, and 3 | 23 | Giusti et al. (2022) |
| 63 | Manzanillas-Castro and Acosta-López | 2022 | Biodiversity (English) | Sharks | Multiplex PCR-SSP | ITS2 | Commercial (fish market in Ecuador) | ECUADOR | 1 and 2 | — | Manzanillas-Castro and Acosta-López (2022) |
| 64 | Marín et al. | 2022 | Marine and Fisheries Sciences (English) | Fish | DNA sequencing | COI, Control Region | Commercial (landings in Peru) | PERU | 1, 2, and 3 | 96 | Marín et al. (2022) |
| 65 | Villate-Moreno et al. | 2022 | PeerJ (English) | Sharks | DNA sequencing | NAD2 | Commercial (landings in Colombia) | COLOMBIA | 1 and 2 | — | Villate-Moreno et al. (2022) |
| 66 | Wenne et al. | 2022 | Aquaculture (English) | Mussels | MassARRAY SNP Genotyping | M7 | Commercial (market in Poland, imported from Chile) | POLAND | 1 and 2 | — | Wenne et al. (2022) |
| 67 | Alfaro-Cordova et al. | 2023 | Conservation Genetics Resources (English) | Devil rays | DNA sequencing | COI | Commercial (landings in Peru) | PERU | 1 and 2 | — | Alfaro-Cordova et al. (2023) |
| 68 | Cardeñosa et al. | 2023 | Animal Conservation (English) | Sharks fins | qPCR-SSP, DNA sequencing | ITS2, COI | Commercial (shark fin seizure at El Dorado International Airport-Colombia) | COLOMBIA | 1, 2, and 3 | 194 | Cardeñosa et al. (2023) |
| 69 | Vera et al. | 2023 | Revista Peruana de Biología (Spanish) | Jewelfish | DNA sequencing | COI | Commercial (landings in Peru) | PERU | 1 and 2 | — | Vera et al. (2023) |
| 70 | Ehemann et al. | 2024a | Fish Biology (English) | *Urotrygon* rays | DNA sequencing | COI, NAD2 | Commercial (landings in Ecuador and Peru) | MEXICO | 1 and 2 | — | Ehemann et al. (2024a) |
| 71 | Ehemann et al. | 2024b | Taxonomy (English) | *Gymnura* rays | DNA sequencing | COI | Commercial (landings in Peru) | MEXICO/PERU | 1 and 2 | — | Ehemann et al. (2024b) |
| 72 | Gil et al. | 2024 | Journal of the Science of Food and Agriculture (English) | Shrimp/prawns | DNA sequencing | COI | Commercial (supermarket and fishmongers in Spain, imported from Ecuador) | SPAIN | 1, 2, and 3 | 12 | Gil et al. (2024) |
| 73 | Ortiz-Moriano et al. | 2024 | Food Control (English) | Shrimp/prawns | DNA sequencing | COI, 16S rRNA | Commercial (supermarkets in Spain, imported from Ecuador) | SPAIN | 1, 2, and 3 | 1 | Ortiz-Moriano et al. (2024) |
| 74 | Tiktak et al. | 2024 | Plos One (English) | Finfish, sharks, rays | DNA sequencing, LOC-LAMP | COI, ITS2, Control Region | Commercial (fishing ports and markets in Ecuador) | UK/ECUADOR | 1 and 2 | — | Tiktak et al. (2024) |
| 75 | Marín et al. | 2025 | Plos One (English) | Shellfish/fish | DNA sequencing, qPCR-SSP, PCR-SSP, multiplex PCR | COI, 16S rRNA, 12S rRNA | Commercial and wild (landings, wholesale, markets, supermarkets in Peru) | PERU | 1, 2, and 3 | 15 | Marín et al. (2025) |

### 3.2 DNA-based identification methods for seafood species from the ESPO region

#### 3.2.1 PCR-RFLP

This technique involves three main steps: 1) the PCR amplification of a specific DNA fragment using universal or group-specific primer sets; 2) enzymatic digestion with one or more restriction endonucleases that recognize specific sites; and 3) the separation of the digested DNA fragments in a gel matrix. A successful RFLP assay will result in the amplicon being cut into two or more fragments, generating species-specific band patterns (Lago et al., 2014). The main advantage of this method relies on its simplicity and relatively low cost. However, one potential drawback is that incomplete enzymatic digestion can lead to misleading diagnostic results, which may necessitate longer incubation times to achieve more reliable outcomes, thereby extending the overall duration of the identification procedure. Additionally, prior knowledge of the target DNA sequence is essential to select the adequate enzyme capable of producing distinct fragment patterns between two or more species (Bossier, 1999; Lago et al., 2014).

PCR-RFLP has been widely utilized to authenticate seafood products globally since the early 90’s (Chow & Inoue, 1993; Chow et al., 1993). In the ESPO region, various commercial species targeted for the development of RFLP assays have been documented in 13 studies. These studies include Chilean mollusks like mussels focusing on various markers such as 18S rDNA, ITS, PAPM, COI, and 16S rRNA (Toro, 1998; Toro et al., 2005; Fernández-Tajes et al., 2011; Jilberto et al., 2017; Larraín et al., 2019; Colihueque et al., 2020; Quintrel et al., 2021; Giusti et al., 2022) and keyhole limpets using the mitochondrial cytochrome b (CYTB) gene (Olivares-Paz et al., 2006). Additionally, PCR-RFLP was used to identify different fish species including canned sardines from Peru based on the CYTB gene (Jérôme et al., 2003a), Chilean hake using the CYTB and 5S rDNA genes (Perez & Vazquez-García, 2004), as well as bream, salmon, and trout from Chile (Espiñeira et al., 2009b) and salmon from Chilean restaurants (Prida et al., 2020) using the CYTB gene.

#### 3.2.2 PCR-RAPD

This approach utilizes short oligonucleotides of arbitrarily chosen sequence (10-12 nt) to generate species-specific electrophoretic patterns, known as fingerprinting, based on the random PCR amplification of several anonymous DNA fragments across the genome (Yamashita et al., 2024). The primary advantage of this method is that it requires little prior knowledge of the target DNA sequence. However, despite its speed and simplicity, the reproducibility of PCR-RAPD can be affected by slight changes in the quality and quantity of the target DNA (Bossier, 1999; Yamashita et al., 2024). As a result, the number of studies reporting RAPD-based protocols has declined over time. Nevertheless, intra- and inter-laboratory reproducible RAPD results can be achieved if reactions are performed under strictly identical conditions (Atienzar et al., 2006).

PCR-RAPD has been applied to species of economic significance from the ESPO, such as differentiating the Mediterranean mussel *Mytilus galloprovincialis* from other commercially valuable mussel species, including the Chilean mussel *M. chilensis* collected from a harvested natural population in Puerto Aguirre, Chile (Rego et al., 2002). In another study, Marín et al. (2007) developed a RAPD-based method for identifying the Pacific abalone *Haliotis discus hannai* and the red abalone *H. rufescens*, as well as their potential hybrids, obtained from a Chilean research institutés hatchery. Based on their RAPD results, the authors reported that the supposed hybrids were actually phenotypic variants of *H. rufescens*.

#### 3.2.3 PCR-SSCP

This method relies on the electrophoretic mobility of single-stranded DNA through a non-denaturing gel. Specific nucleotide mutations present in a single-stranded PCR product alter the moleculés conformation, causing two DNA strands that differ by a single nucleotide to migrate differently (Gasser et al., 2006; Lago et al., 2014). While this method is relatively simple, cost-effective, and time-efficient, it has some disadvantages, such as poor reproducibility and the need for reference samples (Gasser et al., 2006; Silva & Hellberg, 2021). Two studies have demonstrated the application of the PCR-SSCP method to identify fish and abalone species of global commercial interest, including samples collected from the ESPO region. Rehbein et al. (2002) evaluated the utility of PCR-SSCP and RFLP-SCCP assays using a 464 bp fragment of the CYTB gene to identify various fish species in commercially produced fishmeal samples. These samples included anchovy, mackerel, sardine, and hake from Peru and Chile. In another study, Aranceta-Garza et al. (2011) developed a PCR-SSCP technique based on two PCR products targeting partial fragments of the 18S rDNA and Lysin precursor genes in three abalone (Haliotidae) species namely *H. corrugata*, *H. fulgens*, and *H. rufescens*, as well as other gastropod species in raw, frozen, and canned presentations from different origins (Chile, Mexico, and Peru). By utilizing the PCR-SSCP method for the 18S rDNA gene fragment, the authors revealed that a canned abalone sample had been substituted with *Concholepas concholepas* (Muricidae) from Peru.

#### 3.2.4 PCR-SSP

Species identification using species-specific primers (SSPs) is a rapid, highly accurate, and sensitive method. It relies on a perfect or near perfect complementarity between the SSP and the target species’ strand, along with the presence of punctual mutations within the primer hybridization region of non-target species (Marín et al., 2025). Mismatches at the primer’s 3’-terminal of the primer can severely disrupt polymerase activity, leading to the most significant detrimental effects (Stadhouders et al., 2010; So et al., 2020). Several techniques for species identification utilize SSPs, including multiplex PCR, qPCR, eDNA monitoring, and more recently, CRISPR-based methods (Goldberg et al., 2016; Williams et al., 2019; Silva & Hellberg, 2021). The PCR-SSP technique for species identification can be conducted with basic molecular laboratory equipment, and the final diagnostic is electrophoretically resolved based on the presence or absence of a PCR product of a known size. This makes the method both time-effective and easy-to-interpret. Additionally, modified SSPs can be combined with isothermal amplification technologies, using nanoparticle-based colorimetric or lateral flow strip approaches, to enable faster, *in-situ* identification of seafood (Maggioni et al., 2020; Frigerio et al., 2023, Tiktak et al., 2024). The primary drawback of the SSPs approach is the need for prior DNA sequence information from both target and non-target species.

In the ESPO region, the PCR-SSP method has been mentioned in four articles for identifying shark species through fin samples collected in Chile (Hernández et al., 2008, 2010), as well as from artisanal landings in Colombia (Caballero et al., 2012), and a market in Ecuador (Manzanillas-Castro & Acosta-López et al., 2022). Additionally, the PCR-SSP method was used to differentiate the Peruvian anchoveta, *E. ringens* (Chairi & Rebordinos, 2014), and the Peruvian scallop, *A. purpuratus* (Marín et al., 2013, 2017, 2025). Furthermore, this method was proven effective for the accurate identification of a squid and nine fish species of commercial value in the Peruvian seafood sector (Marín et al., 2025).

#### 3.2.5 qPCR-SSP

This method is a faster and more sensitive variant of endpoint PCR-SSP that relies on the real-time detection of a fluorescent signal, which is proportional to the concentration of the DNA product (Silva et al., 2021). Fluorescent chemistries can be derived from non-specific dsDNA-binding dyes (e.g. SYBR Green I) or sequence-specific hydrolysis probes (e.g. TaqMan) (Tajadini et al., 2014). The former technology is relatively inexpensive and easy to use, while the latter offers better specificity because the fluorescent signal is emitted only from specific amplifications (Tajadini et al., 2014). The qPCR-SSP method generally targets relatively short DNA fragments (less than 300bp) and has proven highly effective in identifying seafood species worldwide (Silva et al., 2021), including fish species of the ESPO region, as reported in five articles compiled in this review.

Herrero et al. (2011) successfully developed and applied TaqMan qPCR-SSP assays targeting commercial samples of Atlantic salmon (*Salmo salar*) collected from various countries, including supermarkets in Chile. In another study, Mora et al. (2018) developed novel qPCR-SSP assays for authenticating commercial Peruvian weakfish (*Cynoscion analis*) and hake products (*Merluccius gayi* and *M. peruanus*) based on partial fragments of the ATP6 gene and the control region, respectively. Additionally, Marín et al. (2025) used a partial fragment of the mitochondrial 16S rRNA gene to create a novel qPCR-SSP protocol aimed at authenticating Peruvian scallop samples collected from markets, supermarkets, and restaurants in Peru. qPCR-SSP protocols have also been developed to monitor the trade of ESPO species used in traditional Chinese medicine. Marín et al. (2021) developed a qPCR-SSP assay designed to target a 136 bp fragment of the mitochondrial 16S rRNA gene of the protected Peruvian seahorse, *Hippocampus ingens*. They assessed its effectiveness using specimens confiscated by Peruvian authorities, as well as dried and powdered presentations of seahorses. In a separate study, Cardeñosa et al. (2023) applied a previously established qPCR-SSP protocol (Cardeñosa et al., 2018) to identify CITES-listed sharks using fins that were seized by Colombian authorities that were declared as “dried fish bladders”.

#### 3.2.6 Multiplex-PCR and qPCR

Multiplex-PCR involves the simultaneous amplification of two or more target loci by including multiple primer sets in a single PCR reaction (Markoulatos et al., 2002). This method is both time-saving and cost-effective, allowing for the identification of different species in one run. The result is the generation of species-specific DNA fragments of varying sizes (Rasmussen & Morrissey, 2008), which can be resolved on an agarose gel following endpoint PCR or analyzed through melting curve analyses based on species-specific melt profiles after qPCR (Winder et al., 2011). Some of the disadvantages of this method include the need for proper primer design, which is crucial to avoid the formation of primer dimers and non-specific PCR products, which can deplete the reaction components (Edwards and Gibbs, 1994) and negatively affect efficiency. Additionally, all primers should be designed to have the same or closely related annealing temperatures to ensure effective co-amplification of all target loci. Achieving optimal co-amplification results also requires careful titration of component concentrations, including primers, MgCl2, and dNTPs (Marín et al., 2025), which can be a time-demanding process during PCR reaction standardization.

A total of nine studies reviewed herein assessed the effectiveness of multiplex PCR or qPCR methods for authenticating species from the ESPO region, specifically focusing on sharks (Hernández et al., 2008, 2010; Caballero et al., 2012; Manzanillas-Castro & Acosta-López, 2022; Cardeñosa et al., 2023), and scallops (Marín et al., 2013, 2017). Additionally, five commercially important finfish species from the ESPO region were targeted for species identification using multiplex PCR-SSP assays. These species include the Peruvian anchoveta, *E. ringens*, which was identified through a SSP assays based on the nuclear 5S rDNA gene (Chairi & Rebordinos, 2014). Also, fresh and cooked commercial samples of two congeneric rock seabass, *Paralabrax callaensis* and *P. humeralis*, were successfully identified using a PCR-SSP assay that targeted a partial fragment of the mitochondrial 16S rRNA gene (Marín et al., 2025). In the same study, the authors reported a second multiplex PCR-SSP assay that targeted a partial region of the mitochondrial COI gene for the simultaneous identification of commercial cooked fish that share the same market name “*ojo de uva*” in Peruvian markets: the mocosa ruff *Schedophilus haedrichi*, and the grape-eye seabass *Hemilutjanus macrophthalmos* (Marín et al., 2025). Conversely, less attention has been given to developing multiplex PCR assays for identifying commercial invertebrate species from the ESPO region. Thus far, only the Peruvian scallop, *A. purpuratus*, has been included in two different multiplex PCR-SSP assays targeting the mitochondrial 16S rRNA gene (Marín et al., 2013, 2017).

#### 3.2.7 qPCR-HRM

High-resolution melting (HRM) is a qPCR-based technique primarily used for genotyping single-nucleotide polymorphisms (SNPs) and for species identification. This method monitors the fluorescence signal shift that occurs during the dissociation of the qPCR product from double-stranded to single-stranded DNA in the presence of a saturating dye as the temperature increases (Kim et al., 2023). Specialized HRM software utilizes this fluorescence data to create sequence-specific peak patterns based on the amplicon sequence and temperature of the melting curves (Warnke et al., 2020). This single-tube reaction method is both time- and cost-effective, and it has proven successful in authenticating fish and shellfish species globally (Silva & Hellberg, 2021).

Nevertheless, within the ESPO region, the HRM approach has only been used for the identification of native, introduced, and hybrid commercial mussel species from Chile, specifically *M. chilensis*, *M. edulis*, and *M. galloprovincialis*. This identification is based on the Polyphenolic Adhesive Protein of Mussels (PAPM) (Jilberto et al., 2007; Asorey et al., 2022), and also on a multilocus approach using SNPs discovered through data mining from ESTs and RADseq (Quintrel et al., 2021).

#### 3.2.8 PCR-Microarrays

PCR-Microarrays, also known as DNA chips, traditionally consists of a glass microscopy slide that has oligonucleotide probes attached. These probes are specifically designed to be complementary to the DNA of target species (Teletchea, 2009). Typically, during the PCR process, products are labeled with fluorophores. This labelling allows for the detection of target DNA after hybridization with the species-specific oligonucleotide probes, following the washing steps (Teletchea, 2009). This method enables the simultaneous identification of hundreds, or even thousands, of species. Several studies from different countries across Asia, Europe, and North America have demonstrated its effectiveness for fish identification (Teletchea, 2009; Silva & Hellberg, 2021).

A recent proof-of-concept study (Kappel et al., 2021) assessed the utility of a DNA microarray assay using 21 crustacean species-specific probes targeting a segment of the 16S rRNA gene. This study focused on the giant tiger prawn, *Penaeus (Penaeus) monodon*, and the Pacific whiteleg shrimp *P. (Litopenaeus) vannamei*. The assay proved successful in identifying commercial samples of *P. (Litopenaeus) vannamei* from Ecuador. Importantly, all probes designed to recognize *P. (Litopenaeus) vannamei* sequences displayed no cross-reactivity with any other crustacean species examined, highlighting the high specificity of the reported microarray assay.

#### 3.2.9 LAMP

LAMP is a fast, cost-effective, and sensitive method that enables DNA amplification at an isothermal temperature without the need for a thermal cycler. It typically requires a set of 6 especially designed primers that recognize 8 different regions of the target sequence, resulting in higher specificity (Sahoo et al., 2016). It utilizes a DNA polymerase with strand displacement activity, which is crucial for loop-mediated isothermal amplification (Yang et al., 2024). The main advantages of this technique are its high specificity, sensitivity, and faster performance compared to traditional PCR. Additionally, it requires basic equipment, making it suitable for field applications. The main disadvantage is that the primer design is more complex than in PCR, and the standardization process can be cumbersome and time-consuming (Silva & Hellberg, 2021).

Tiktak et al. (2024) developed a Lab-on-a-Chip (LOC) systems integrating field-based DNA extraction and LAMP detection of three CITES-listed shark species: *Alopias pelagicus*, *A. supercilious*, and *Isurus oxyrinchus*. These systems target a partial region of the mitochondrial control region in *I. oxyrinchus* and *A. pelagicus*, while the nuclear ITS2 region was used in *A. supercilious*. The authors reported successful and fast (full analyses lasted less than one hour) species-level identifications of different shark samples collected from fishing ports and markets in Ecuador.

#### 3.2.10 FINS and DNA Barcoding

Although both strategies share a similar rationale, the FINS method for species authentication was first introduced in 1992 by Bartlett and Davidson (1992) to identify tuna species using the DNA sequence information from the CYTB gene. Later, in 2003, Paul Hebert and colleagues proposed the concept of DNA barcoding, which focuses on using a partial region of the mitochondrial COI gene to identify species across all animal phyla, except for the Cnidaria (Hebert et al., 2003). Both species identification methods consist of three main steps: 1) PCR amplification of a specific partial DNA fragment from the target sample; 2) Sanger sequencing of the PCR product; and 3) phylogenetic analysis based on genetic distance (Tamura and Nei, 1993) and Neighbor-Joining methods (Saitou and Nei, 1987), in which the query sequence is compared to related reference sequences obtained from databases. The final results are presented in a phylogenetic tree, where the unknown sample clusters with the reference sequences that exhibit the fewest nucleotide substitutions or genetic distance (Bartlett & Davidson, 1992; Rasmussen & Morrissey, 2008). A key difference between FINS and DNA barcoding is that FINS relies on informative sites and phylogenetic relationships derived from vouchered reference organisms (Li et al., 2011). In contrast, DNA barcoding follows a standardized procedure that emphasizes the presence of a “barcode gap”. This gap indicates sufficient interspecific genetic distance to allow for species resolution (Bucklin et al., 2011; Hellberg et al., 2016).

This review indicates that DNA sequencing methods, such as FINS or DNA barcoding, were the most commonly used method for seafood species identification, appearing in 44 (58.7%) out of the 75 papers compiled for this study. For instance, DNA barcoding was utilized to examine mislabeling at different stages of the supply chain in Peru and Chile (Marín et al., 2018; Biffi et al., 2020; Velez-Zuazo et al., 2021; Dufflocq et al., 2022). Additionally, it was employed to uncover unreported fish species in landings from both Peru and Chile (Canales-Aguirre et al., 2018; Marín et al., 2022). Other studies have utilized traditional taxonomic methods combined with DNA barcoding to describe new distribution ranges, determine species boundaries, and document rare occurrences in fish samples from Peruvian artisanal fisheries (Cabanillas-Torpoco et al., 2020; Cañedo-Apolaya et al., 2021; Vera et al., 2023).

Additional economically significant marine species from the ESPO region identified through DNA sequencing include sharks from landings in Peru (Velez-Zuazo et al., 2015) and Colombia (Caballero et al., 2012; Villate-Moreno et al., 2022), Myliobatiform rays collected from artisanal fisheries in Peru and Ecuador (Alfaro-Cordova et al., 2023; Ehemann et al., 2024a), and a variety of crustaceans (crabs and shrimps) and mollusks (clams, scallops, snails, squids, and octopus) from the Peruvian fishery sector (Marín et al., 2018). Furthermore, DNA analysis has been applied to crab meat products in Chilean markets (Haye et al., 2012), cephalopod species from Ecuador (Pliego-Cárdenas et al., 2016), and non-traditional seafood products such as cetacean meat sold in Peruvian markets (Tzika et al., 2010; Biffi et al., 2020). A DNA mini-barcoding approach, which is a variation of the full DNA barcoding, utilizes short DNA sequences (typically shorter than 200 bp) and has been effective in identifying seafood species from highly degraded samples, such as processed seafood products (Fernandes et al., 2021). Marín et al. (2018, 2025) successfully applied this mini-barcoding method, focusing on short fragments of the mitochondrial COI and 12S rRNA genes, to authenticate various cooked fish samples sourced from Peruvian restaurants. It is important to note that none of these samples were amplifiable using different full DNA barcoding markers.

In the addition to the COI marker, numerous studies have reported the utility of DNA sequences from other mitochondrial gene regions for seafood authentication, such as the 16S rRNA gene. This genetic marker was utilized by Ardura et al. (2012) and Horreo et al. (2013) to authenticate fish feed samples from various origins, including the Peruvian anchovy, *E. ringens*, which is marketed in France, Germany, and Spain. Similar, the same marker was also employed to authenticate scallop samples collected worldwide, including the Peruvian scallop, *A. purpuratus*, sourced from seafood markets in Peru, Germany, and Spain (Näumann et al., 2012; Marín et al., 2015, 2018; Klapper & Schröeder, 2021; Parrondo et al., 2021).

The performance of some nuclear genetic markers has also been assessed in FINS studies aimed at identifying seafood species. For instance, Espiñeira et al. (2009a) highlighted the effectiveness of the ribosomal 18S rDNA gene for identifying commercially important marine bivalves collected from different regions worldwide, including Chile and Peru. Conversely, a study by Asorey et al. (2022) showed that FINS based on the histone H1C gene were not effective for distinguishing between the mussels *M. chilensis* and *M. galloprovincialis* sourced from a growing center in Chile.

#### 3.2.11 HTS NGS-metabarcoding

Next-generation sequencing (NGS) technologies enable the simultaneous sequencing of hundreds of millions of DNA molecules from multiple samples within a relatively short timeframe (Fernandes et al., 2021; Silva & Hellberg, 2021). The NGS approach focused on species identification is known as “NGS metabarcoding”. This method utilizes universal primers to achieve massive amplification of barcode sequences from different species in a single run (Fernandes et al., 2021). Numerous studies reported successful species identification of seafood samples (e.g. tuna, cod, bivalves, clupeids, flatfish) using specific HTS techniques based on Illumina, pyrosequencing, and Ion Torrent platforms (Silva & Hellberg, 2021; Silva et al., 2021).

Among these NGS technologies, 454 pyrosequencing by Roche was the first commercially available platform that used real-time detection of pyrophosphate (PPi) molecules released during DNA sequencing. This process results in light emission from the firefly enzyme luciferase, which is visualized as specific peaks corresponding to individual nucleotides (Fernandes et al., 2021; Silva & Hellberg, 2021). This method has the advantage of providing fast and high accurate performance, and the ability to produce longer reads (Fernandes et al., 2021). Abbadi et al. (2017) utilized 454-pyrosequencing based on the mitochondrial COI and 16S rRNA genes to successfully identify 15 different bivalve species, including both fresh and processed samples of the Chilean mussel, *M. chilensis*, sourced from markets in Italy.

Currently, the Illumina platform is the most widely used NGS technology. This method relies on the PCR amplification and sequencing of DNA clusters that are immobilized on a disposable flow cell, using reversible fluorescent dideoxy terminators (Fernandes et al., 2021; Silva & Hellberg, 2021). One of the major advantage of this approach over traditional Sanger sequencing is its higher throughput per run and lower cost per base (Silva & Hellberg, 2021). Gense et al. (2021) reported the development of a DNA metabarcoding assay using the MiSeq and Iseq Illumina platforms, which focused on a fragment of the mitochondrial 16S rRNA gene. This assay was used to authenticate local and imported oyster, mussel, and scallop seafood products purchased from different commerce in Austria. The authors accurately identified a frozen Peruvian scallop (*A. purpuratus)* sample that was correctly labelled as “*purpur kammmusc*hel” (German for “purple scallop), as well as 17 mussel products that claimed to contain the Chilean mussel, *M. chilensis*. However, these products were identified as two different mussel species: *M. edulis* and *M. galloprovincialis*.

#### 3.2.12 Mass spectrometry-based SNP genotyping

Mass spectrometry is one of the most commonly used method for protein identification and quantification (Cerqueira et al., 2024). However, it can also be applied to the analysis of nucleic acids, including the detection of SNPs and indels (Ellis & Ong, 2017). The process for SNP detection using mass spectrometry involves coupling it with endpoint PCR. In this method, targeted SNPs are amplified through standard PCR, followed by a “single base PCR extension” reaction. This reaction employs mass-modified dideoxynucleotide (ddNTPs) terminators in an oligonucleotide primer (called the extension primer) that anneals just upstream of the polymorphic mutation site. Finally, the mass spectrometer detects the distinct mass of the extended primer within the PCR product, thereby identifying the SNP allele (Gabriel et al., 2009). Wenne et al. (2022) reported the development of a multi-locus SNP assay using mass array spectrometry to authenticate mussel samples. This assay is based on a panel of 54 SNPs from two mussel genes: Me15–16 (a nuclear adhesive protein gene marker) and M7 (an acrosomal sperm protein called M7 lysin). The purpose of this assay was to assess the species and geographical origin information declared in the label of mussel products sold in EU markets. Reference samples from various *Mytilus* species, including farmed *M. chilensis* collected in the province of Chiloe (Chile), were used in this analysis.

### 3.3 Molecular markers and primers

A total of 16 genetic markers were used to identify the ESPO species. Half of these markers originated from the mitochondrial genome, including ATP6, COI, COIII, CYTB, NAD2, 12S rRNA, 16S rRNA, and the Control Region. The other half came from the nuclear genome and included M7, M1C, MAC-1, ITS, PAPM, Rhodopsin, 5S rDNA, and 18S rDNA. The relationships among the molecular markers, target species, and detection methods discussed in the surveyed articles are presented in the form of a Sankey diagram in Fig. 1.

**Fig. 1.**
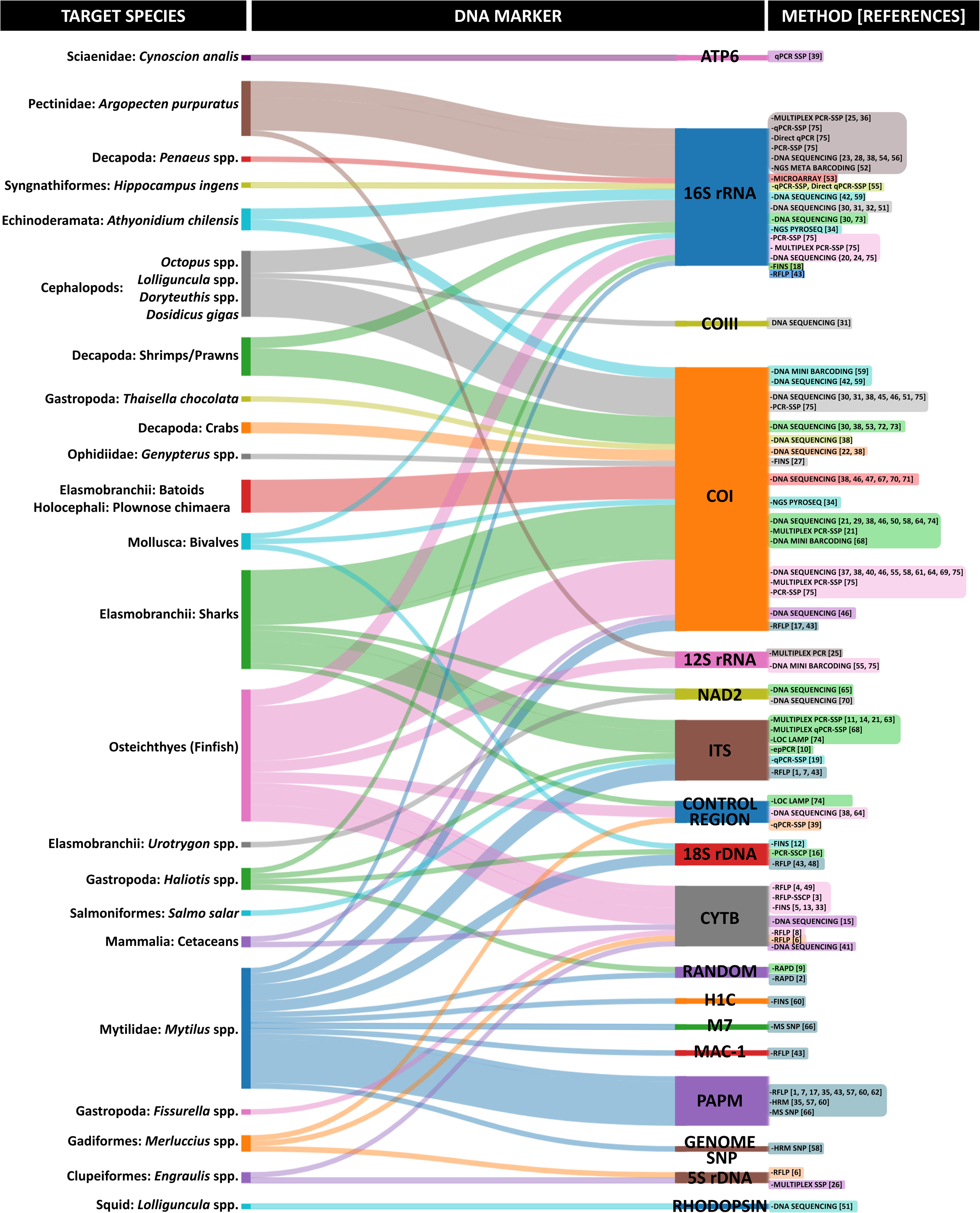
Sankey diagram illustrating the relationships between target seafood species and DNA markers. The width of each connector is proportional to the number of articles reporting the use of each DNA marker. The molecular methods associated with the DNA markers are shown, along with their corresponding article reference numbers (in brackets), which can be found in Table 1.

The mitochondrial COI gene was the most frequently used marker, appearing in 35 articles (46.7%). It was primary applied in DNA sequencing identification methods, including DNA barcoding, mini barcoding, FINS, and pyrosequencing, to identify fish, mollusks, crustaceans, cetaceans, and holothurians. The mitochondrial 16S rRNA gene was the second most commonly utilized marker, featured in 24 articles (32%). This gene was employed in sequencing and species-specific primers detection methods targeting crustaceans, fish, holothurians, and mollusks. The nuclear ITS region and the mitochondrial CYTB gene were the third and fourth most assessed marker, with 11 articles (14.7%) and 10 articles (13.3%) respectively. The nuclear ITS region was utilized in species-specific primer and RFLP detection methods targeting gastropods, mussels, sharks, and salmon. The CYTB gene was used in DNA sequencing and RFLP methods focused on fish and cetaceans.

Most mitochondrial markers reviewed herein have been used to identify multiple animal groups, including the COI, 16S rRNA, 12S rRNA, and CYTB genes. In contrast, four mitochondrial markers were specifically employed to identify individual seafood types: ATP6, NAD2, and the control region for fish, and the COIII for cephalopods. Among nuclear markers, the ITS and the 18S rDNA were used to target multiple seafood types, while the H1C, M7, Mac-1, and PAPM genes were dedicated solely to authenticate mussels. The 5S rDNA was used for fish identification, and Rhodopsin was designated for cephalopods.

The DNA marker assessed using the highest number of detection methods corresponds to the mitochondrial 16S rRNA gene. This marker was utilized in ten different methods including multiplex PCR-SSP, PCR and qPCR-SSP, Direct qPCR-SSP, DNA sequencing (FINS and barcoding), NGS meta-barcoding, microarray, NGS pyrosequencing, and RFLP. These findings demonstrate the high utility and effectiveness of this mitochondrial marker for identifying a wide range of seafood species. This effectiveness is supported by the broadly targeted region spanned by the universal primers developed by Palumbi (1991), as well as by the extensive number of available 16S rRNA gene sequences in public databases.

A descriptive analysis of the primers used in the reviewed articles is provided in Supplementary File S6. This section includes a detailed list of all the primers identified in the articles, along with their associated molecular markers, targeted species, and original reference sources.

### 3.4 Species diversity

A total of 204 species from the ESPO were identified by DNA-based methods reported in the articles reviewed herein. These species were distributed among 142 genera, 82 families, and 44 orders (Table 2, Fig. 2). Fish accounted for the majority of the identified samples (152 species, 74.5%), followed by mollusks (40 species, 19.6%), crustaceans (7 species, 3.4%), cetaceans (4 species, 2%), and holothurians (1 species, 0.5%). Out of the total species identified, 192 were native to the ESPO, whereas 10 were introduced species. The introduced organisms included 6 fish species (*Oncorhynchus kisutch*, *O. mykiss*, *O. tshawytscha*, *Salmo salar, S. trutta*, and *Rachycentron canadum*), two mussel species (*Mytilus edulis* and *M. galloprovincialis*), and two abalone species (*Haliotis discus hannai* and *H. rufescens*).

**Fig. 2.**
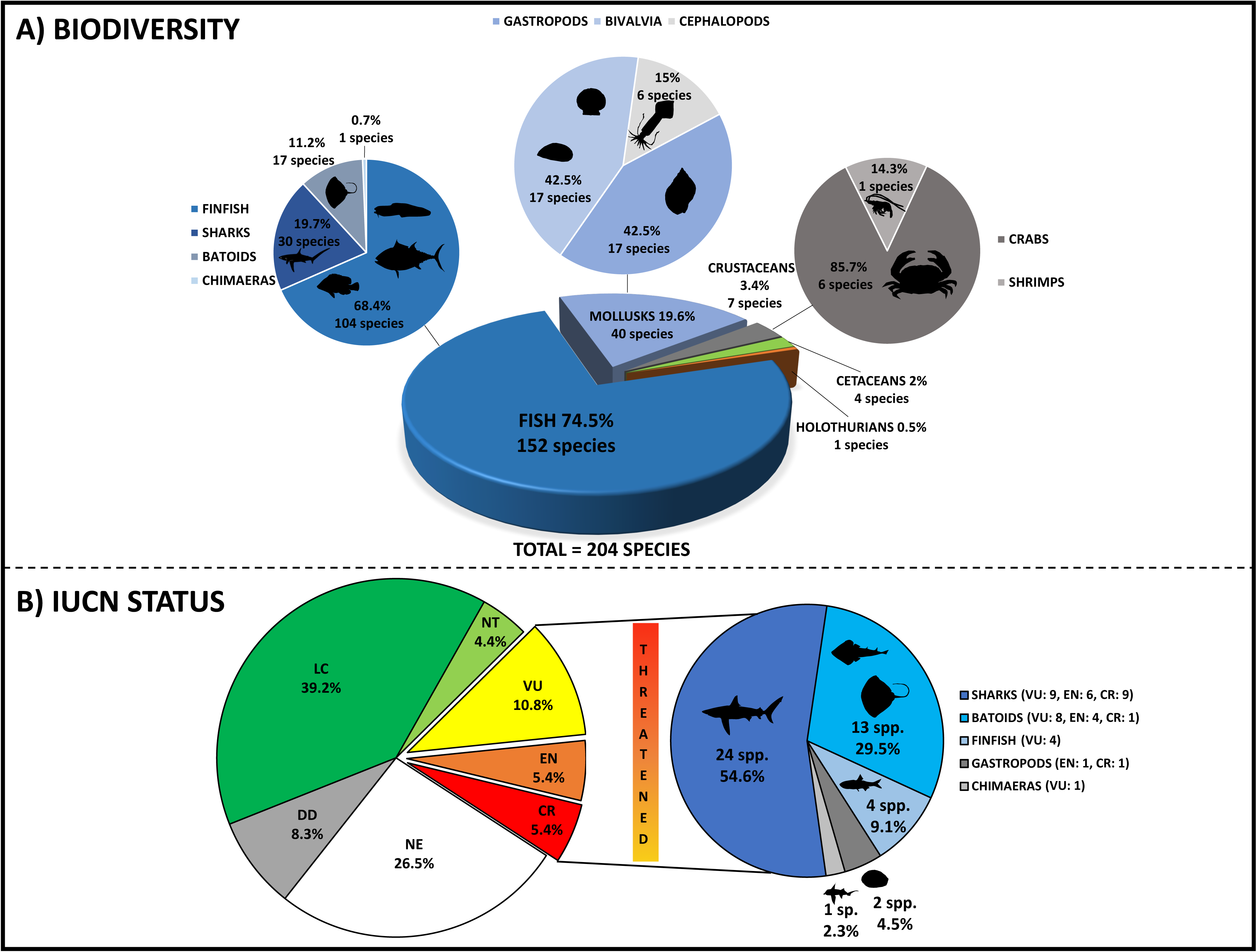
Pie charts in Panel A show the taxonomic diversity of the seafood species identified using DNA-based methods. Species detected were grouped into five major taxonomic categories: fish (blue), mollusks (pale blue), crustaceans (dark grey), cetaceans (green), and holothurians (orange). Wedge sizes represent the proportion of identified species assigned to each category, based on the number of species identified across all reviewed studies. Panel B shows a pie chart illustrating the proportion of total species, which are categorized by their conservation statuses, according to data from the IUCN. The satellite pie chart shows the proportions and taxonomic diversity of threatened species (Vulnerable: VU, Endangered: EN, and Critically Endangered: CR). These include sharks (blue), batoids (deep sky blue), finfish (pale blue), gastropods (dark grey), and chimaeras (light grey).

**Table 2.**
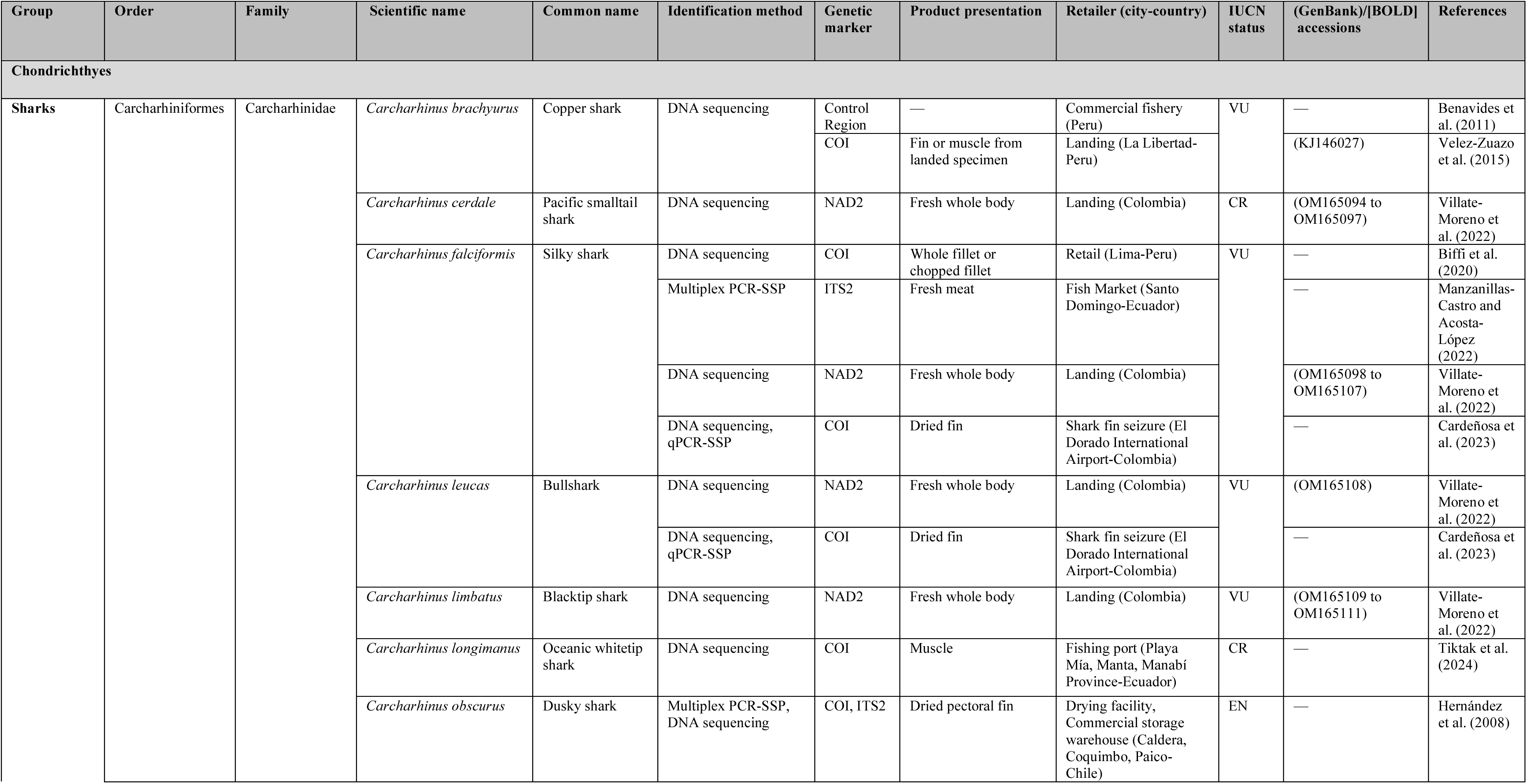
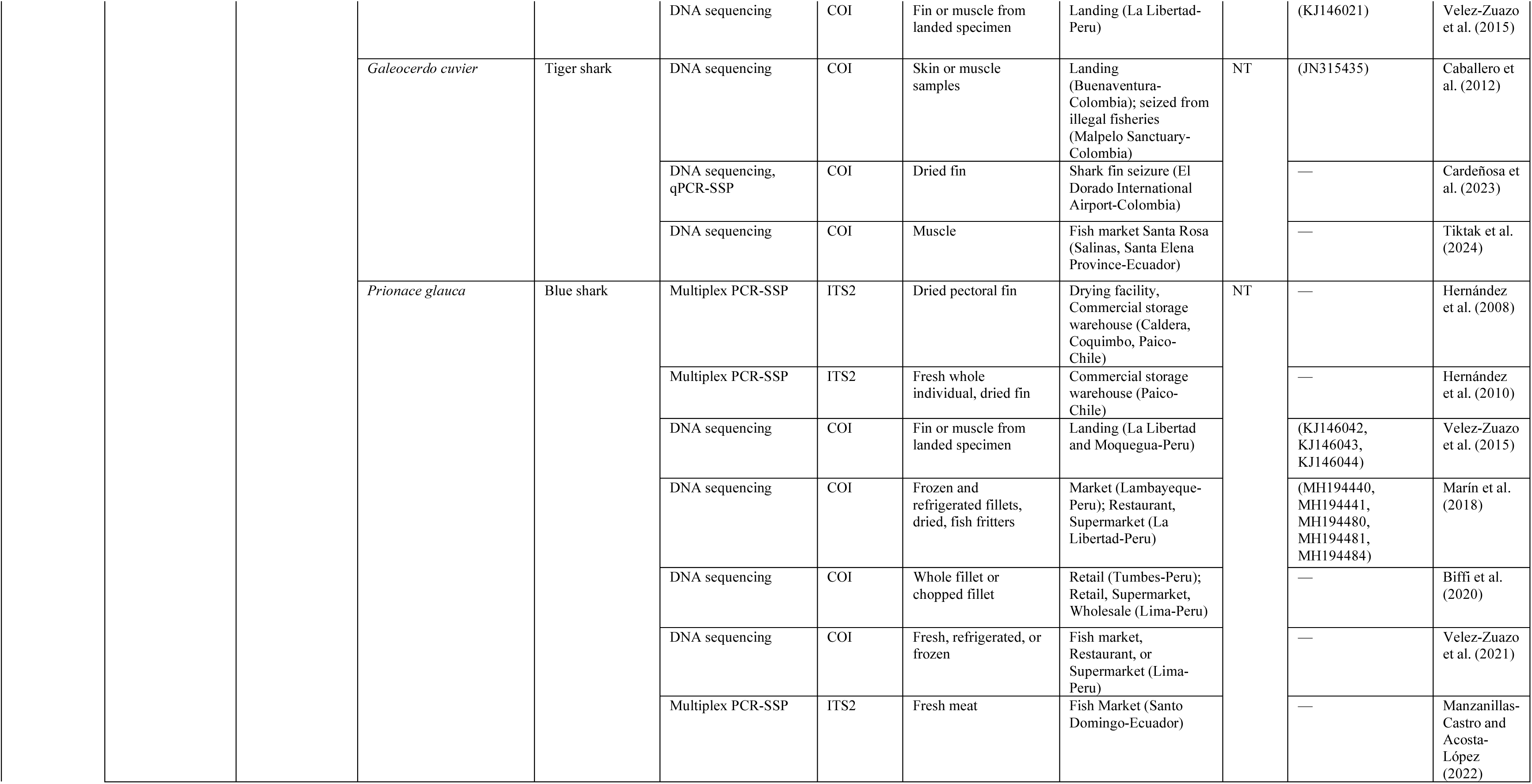
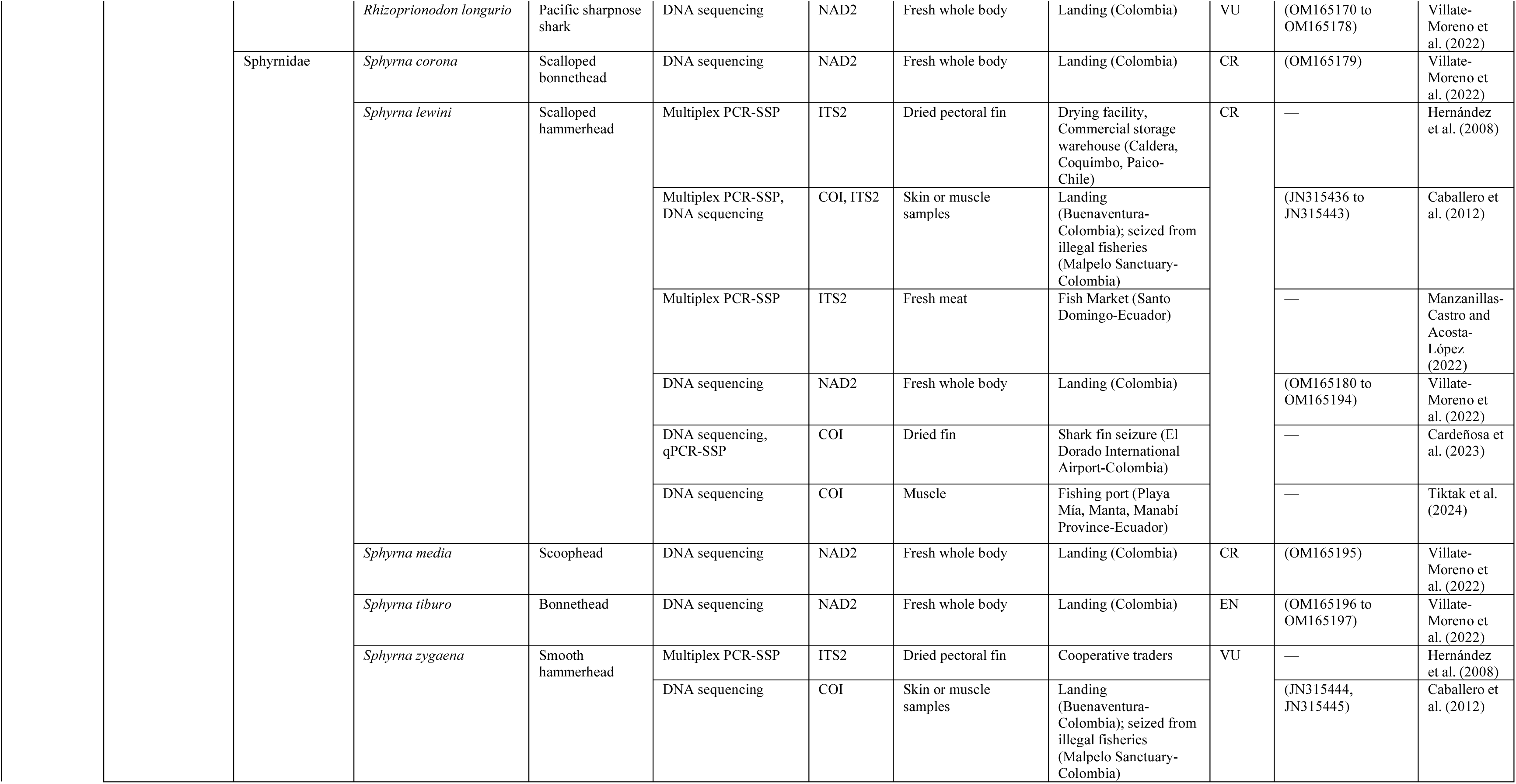
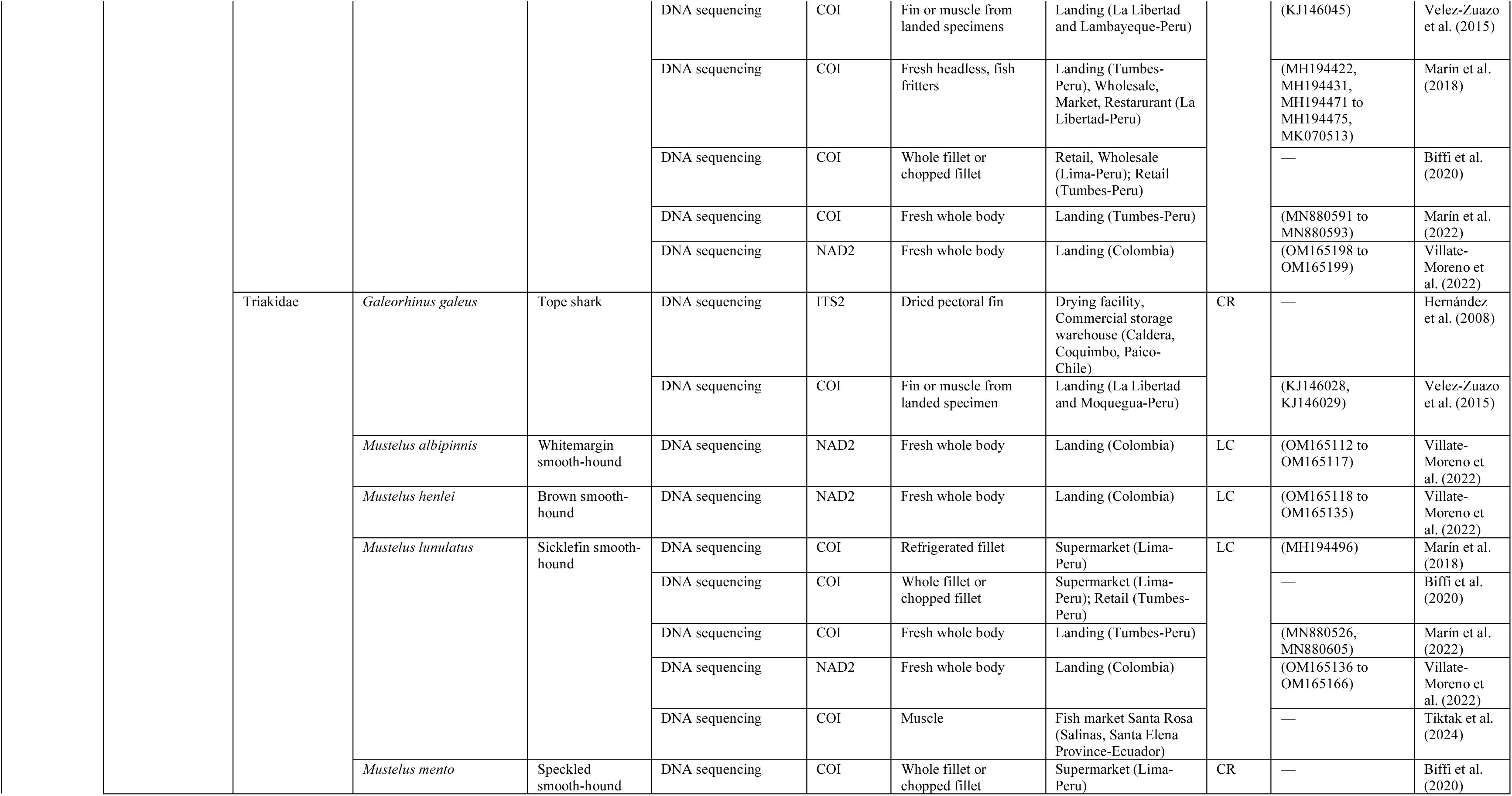
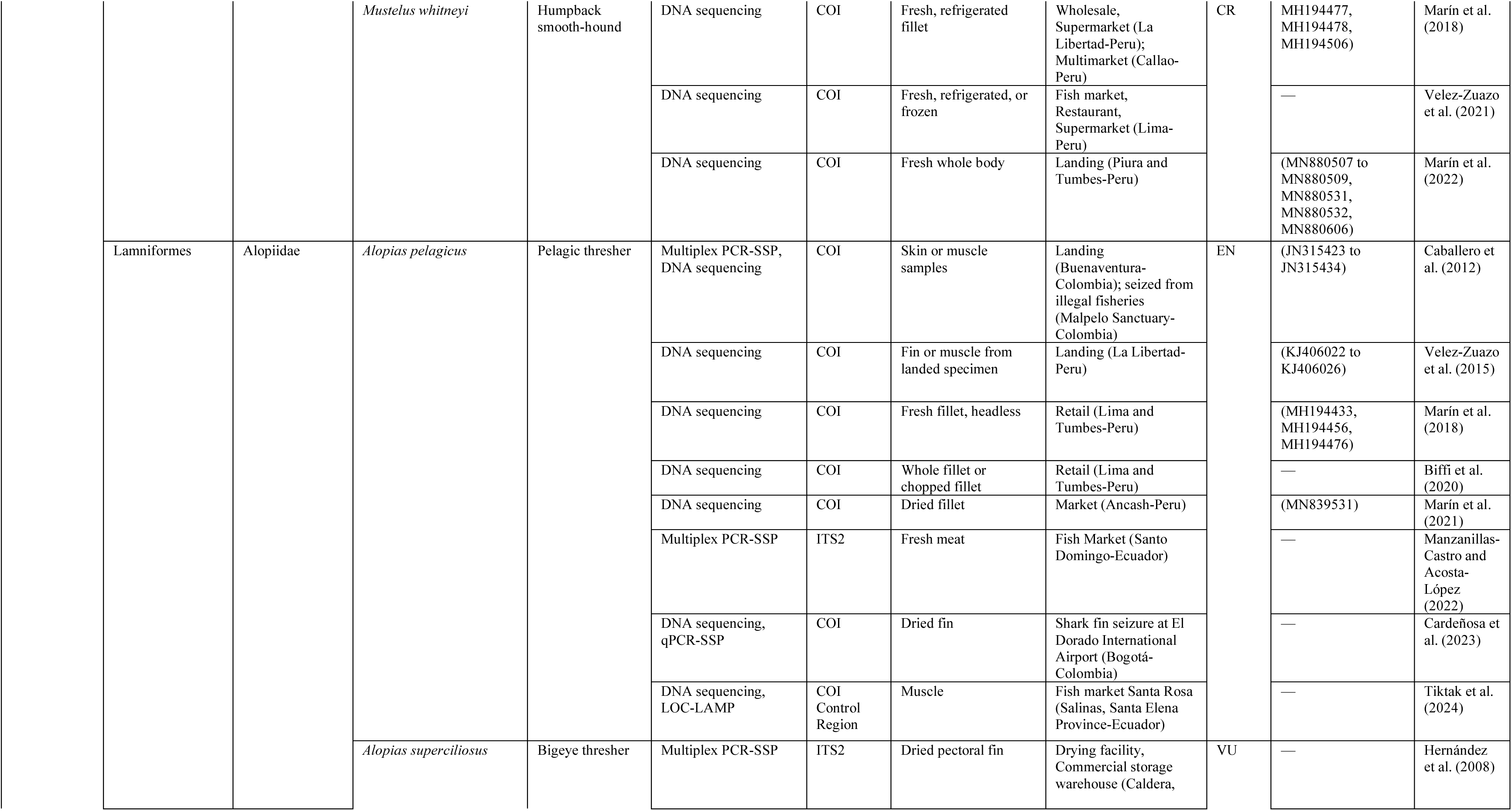
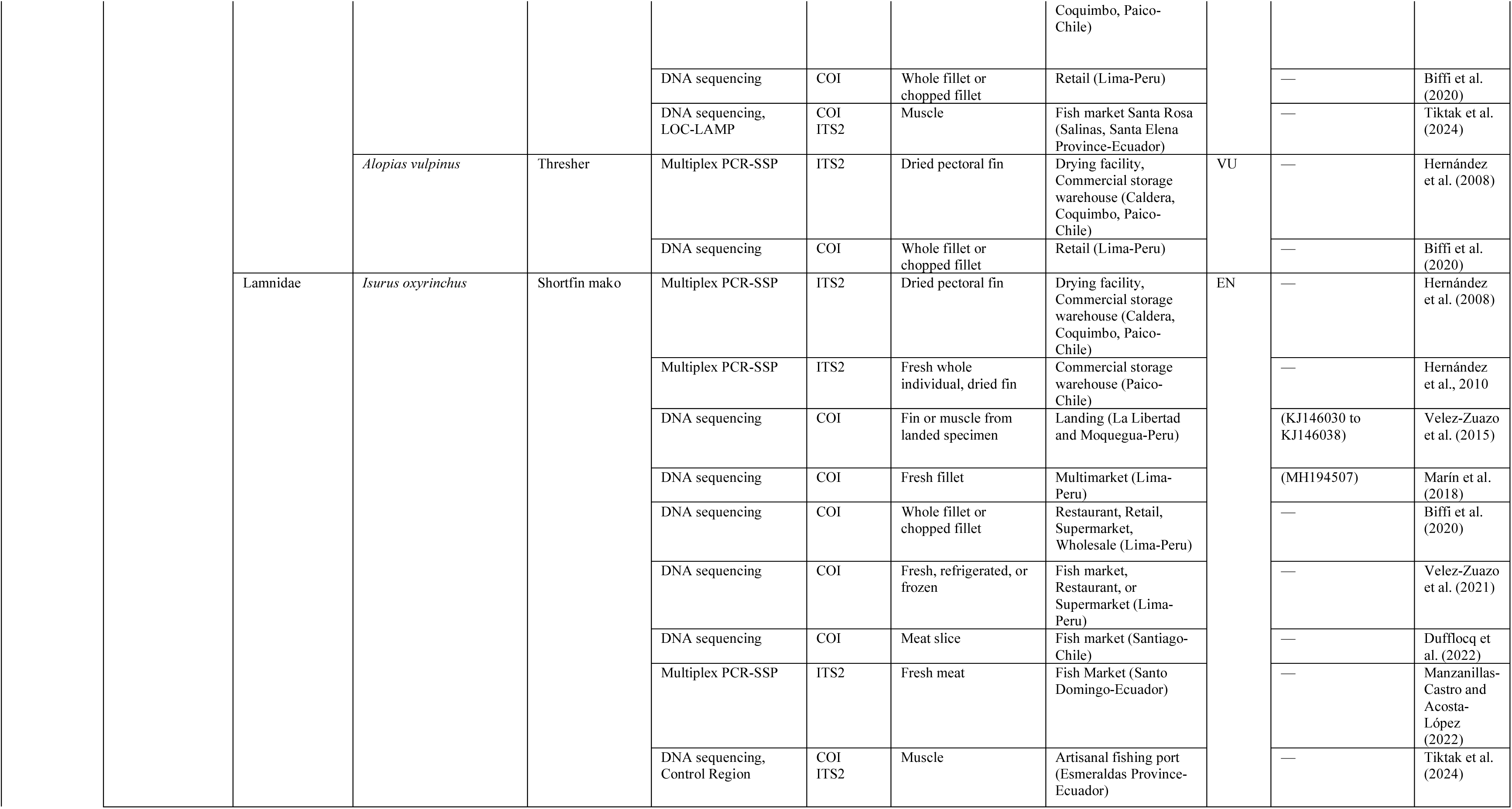
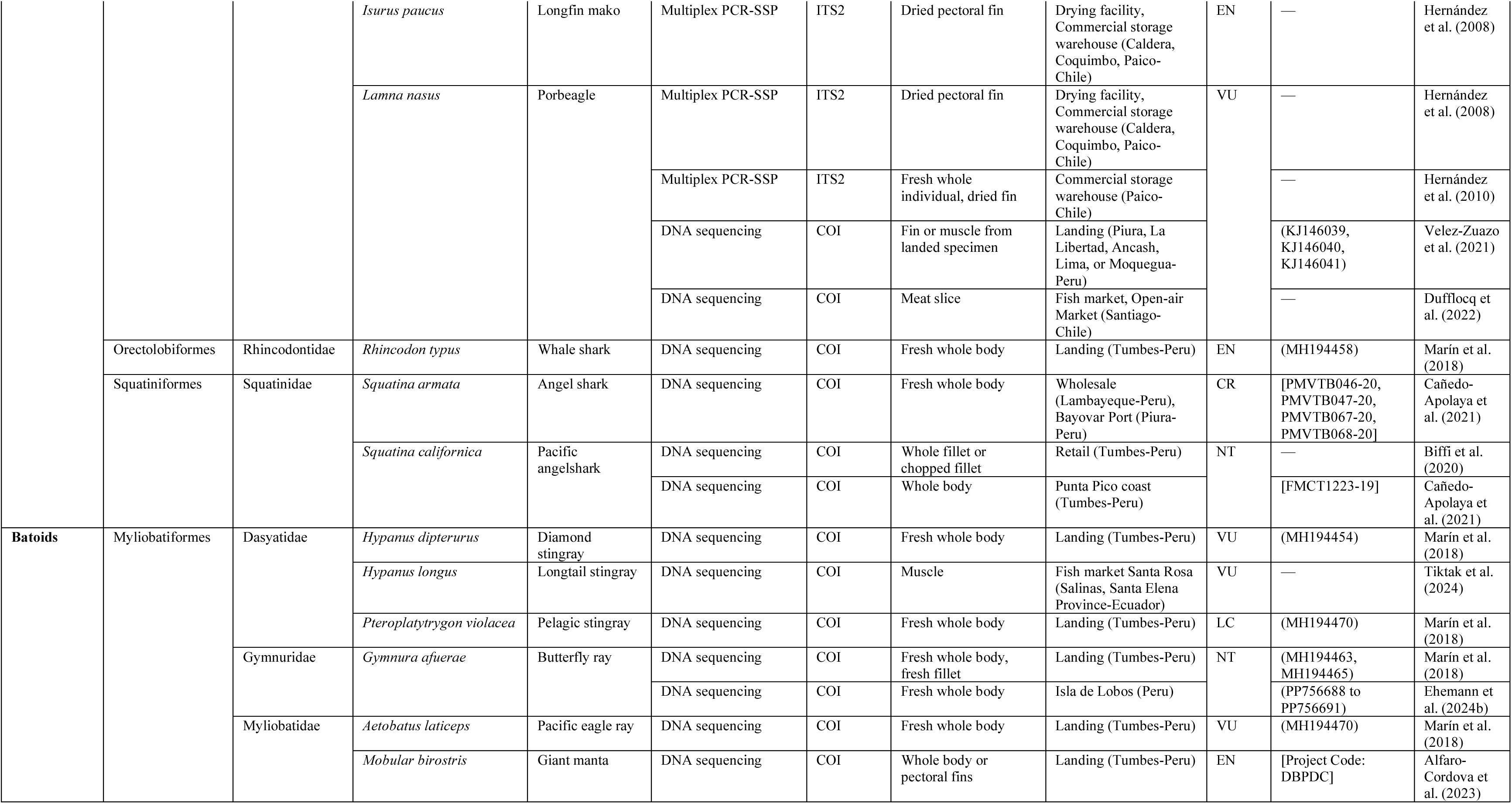
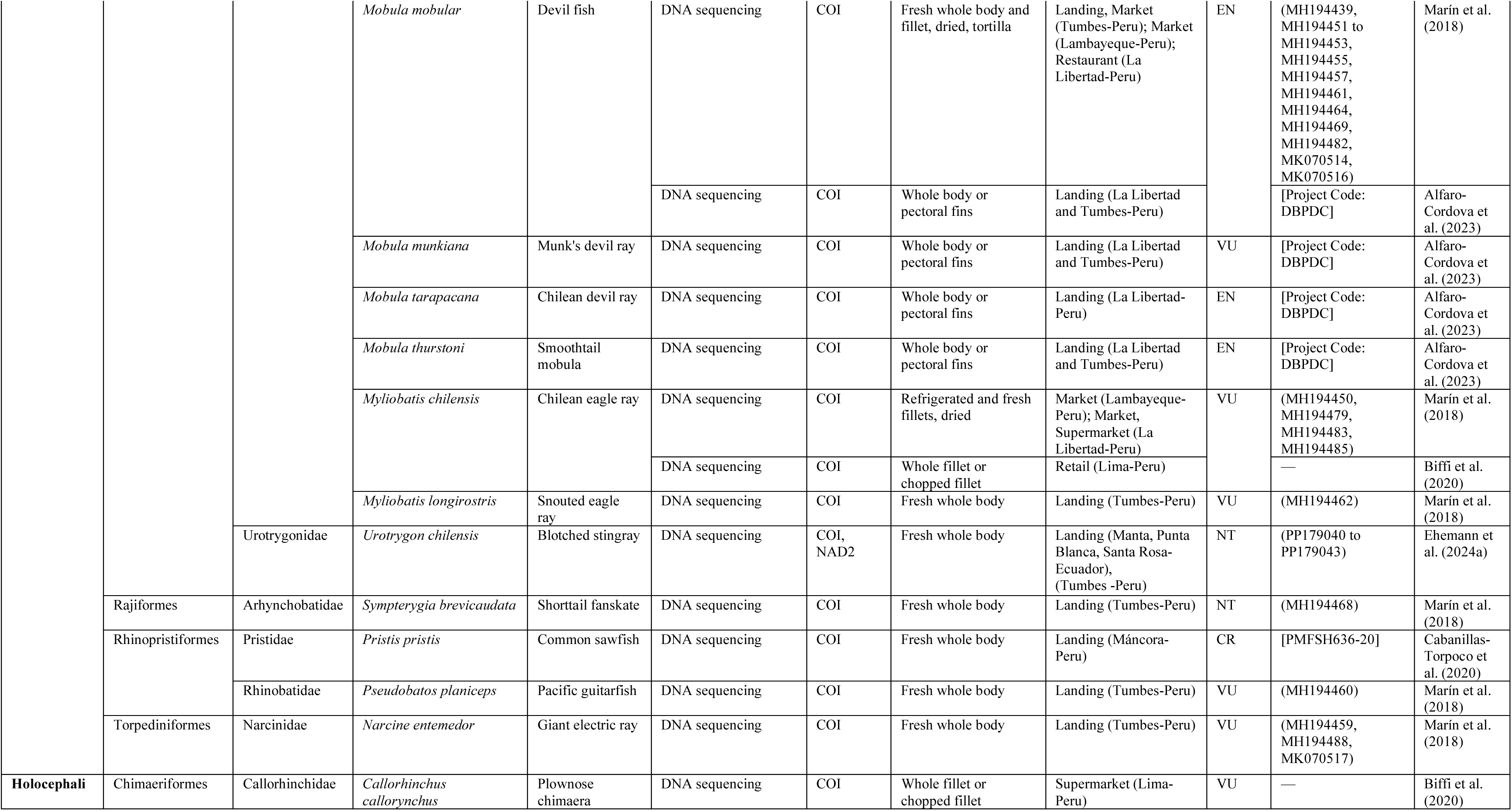
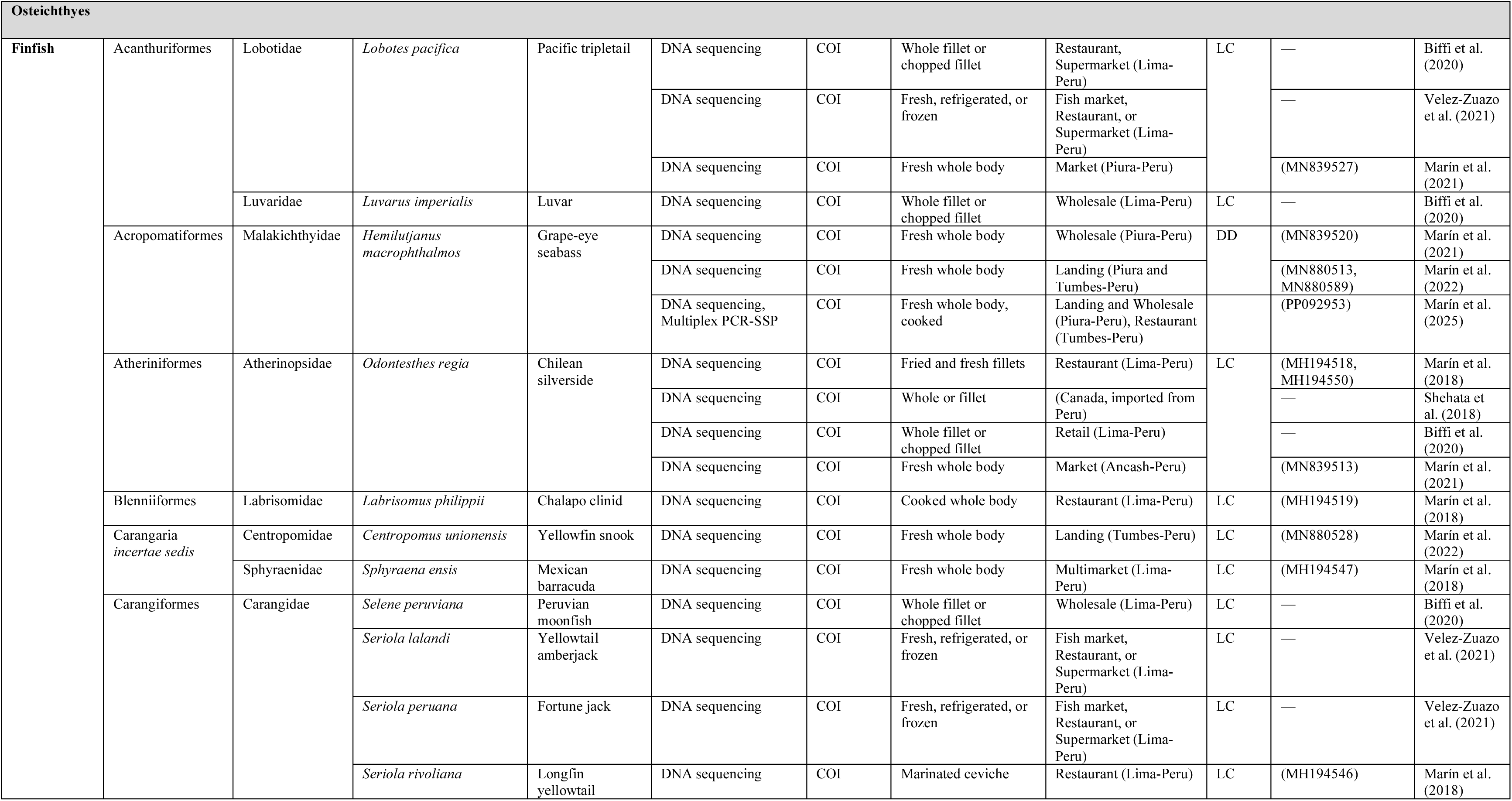
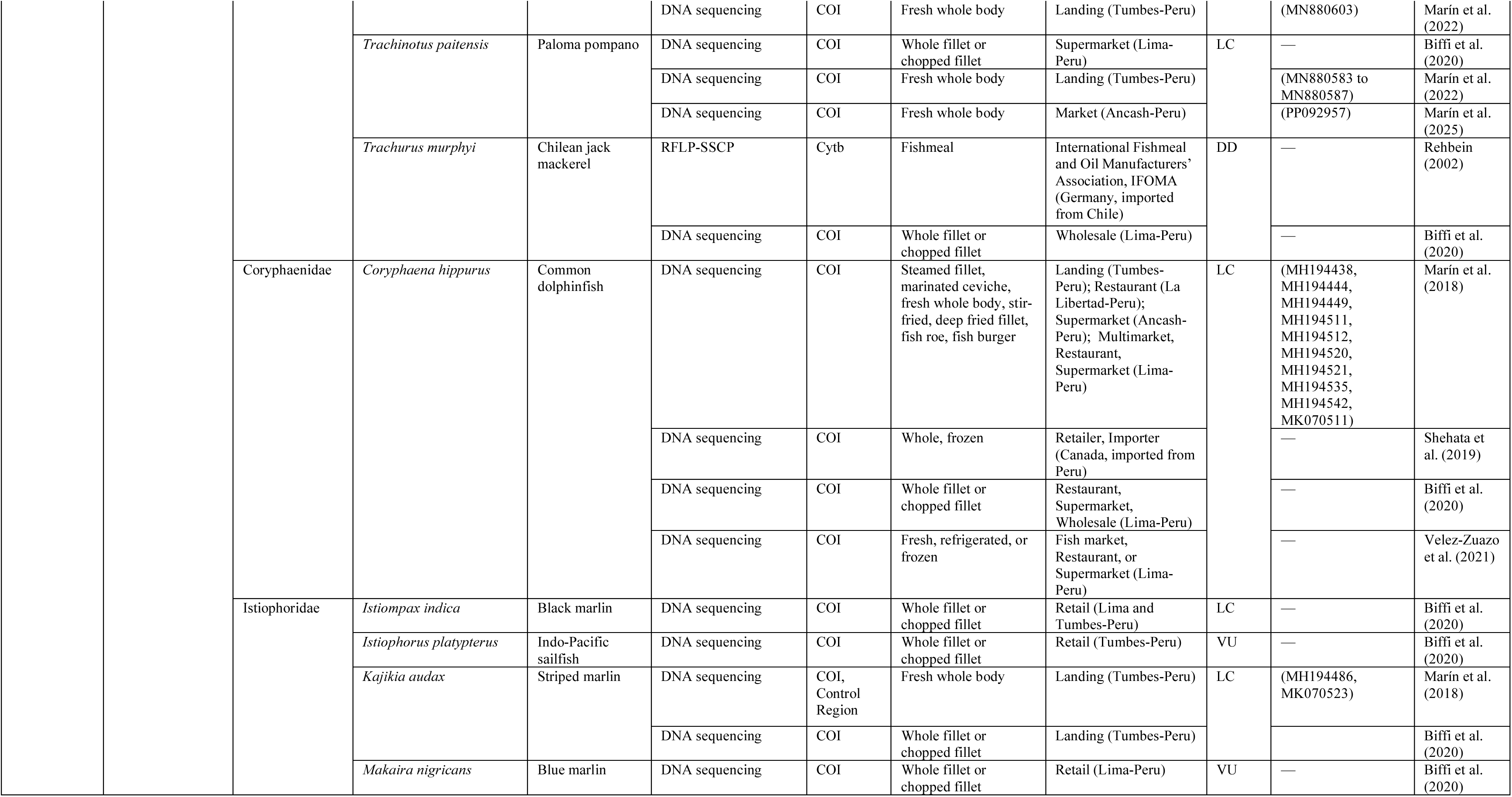
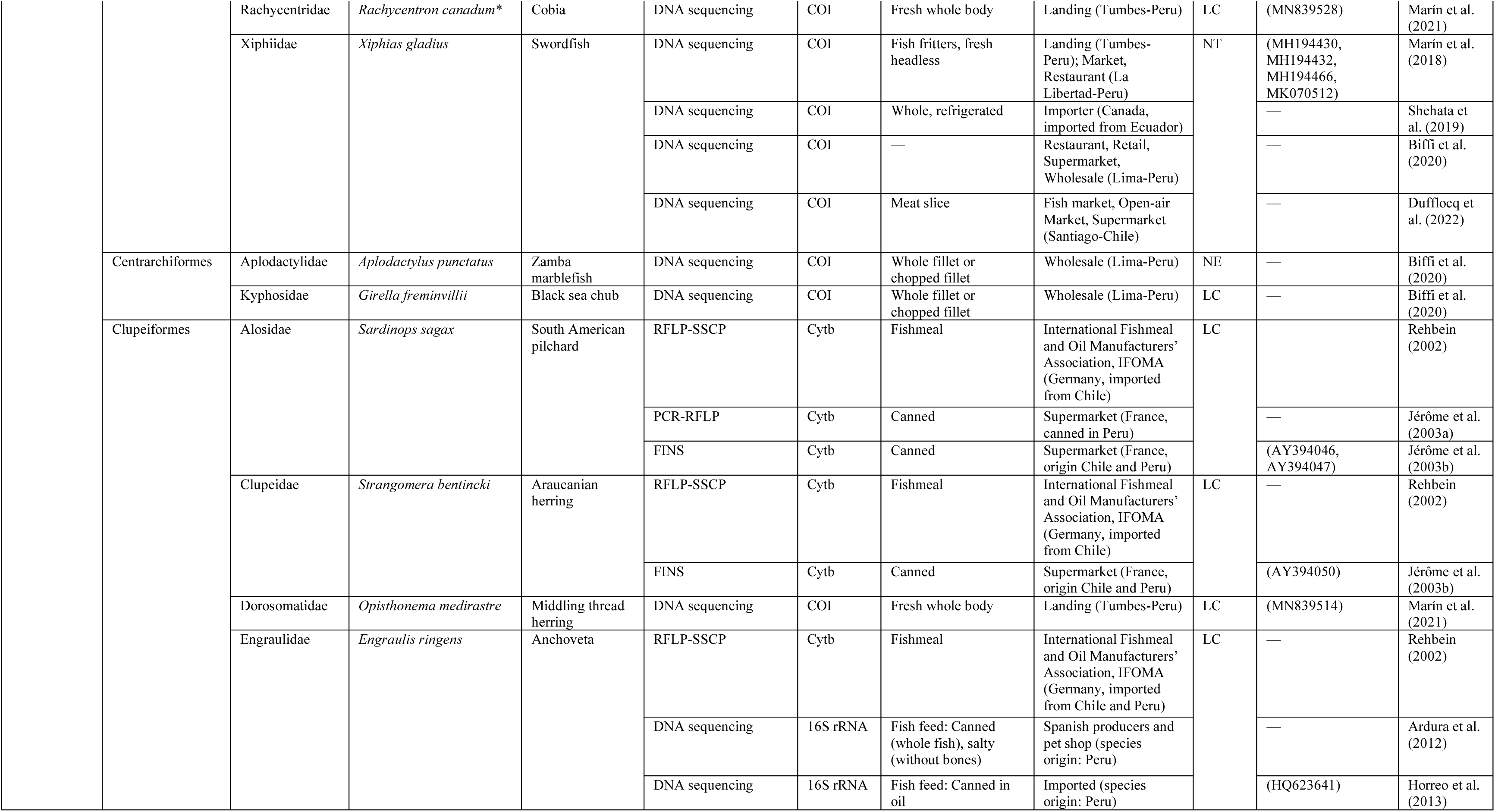
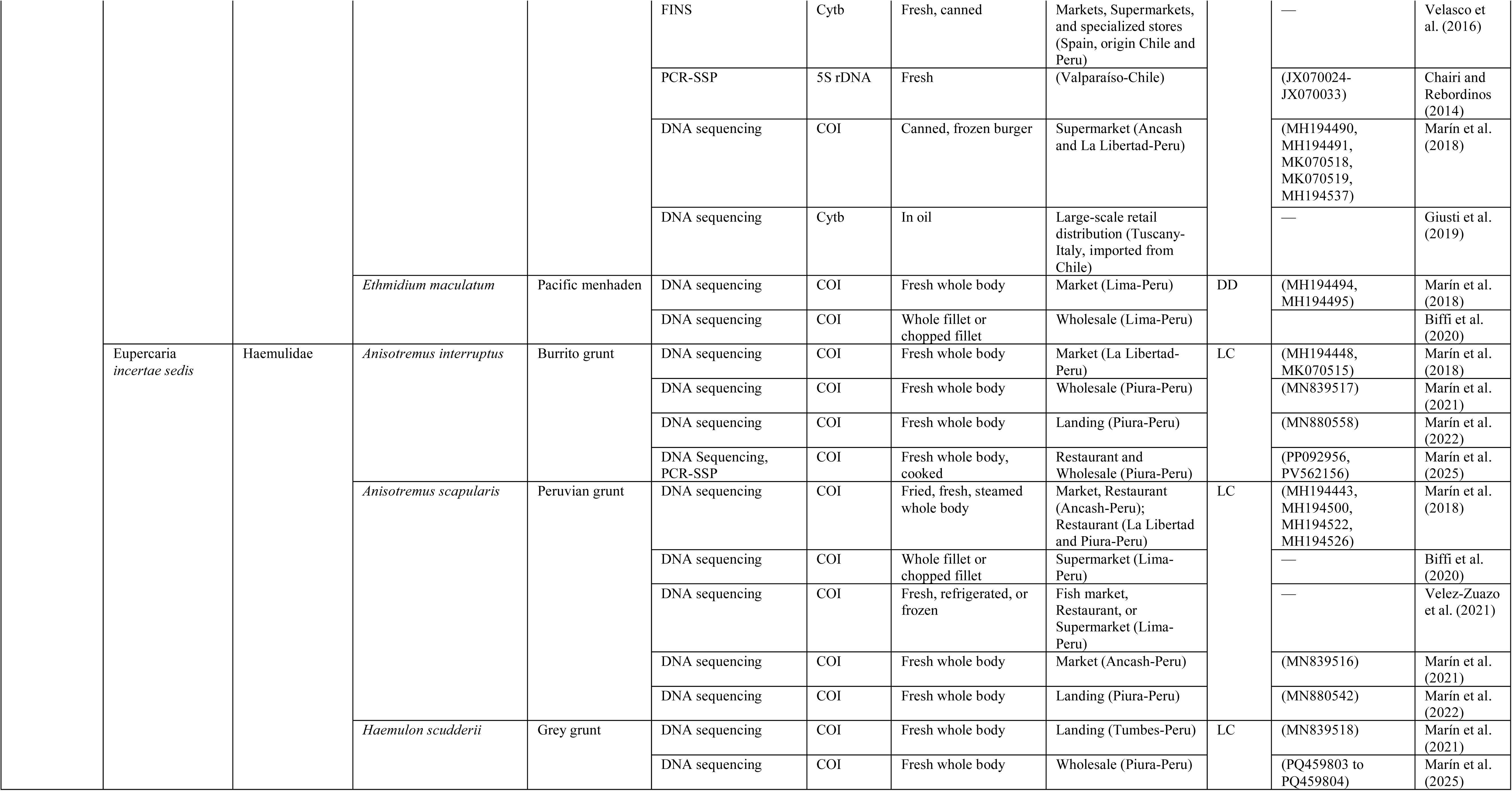
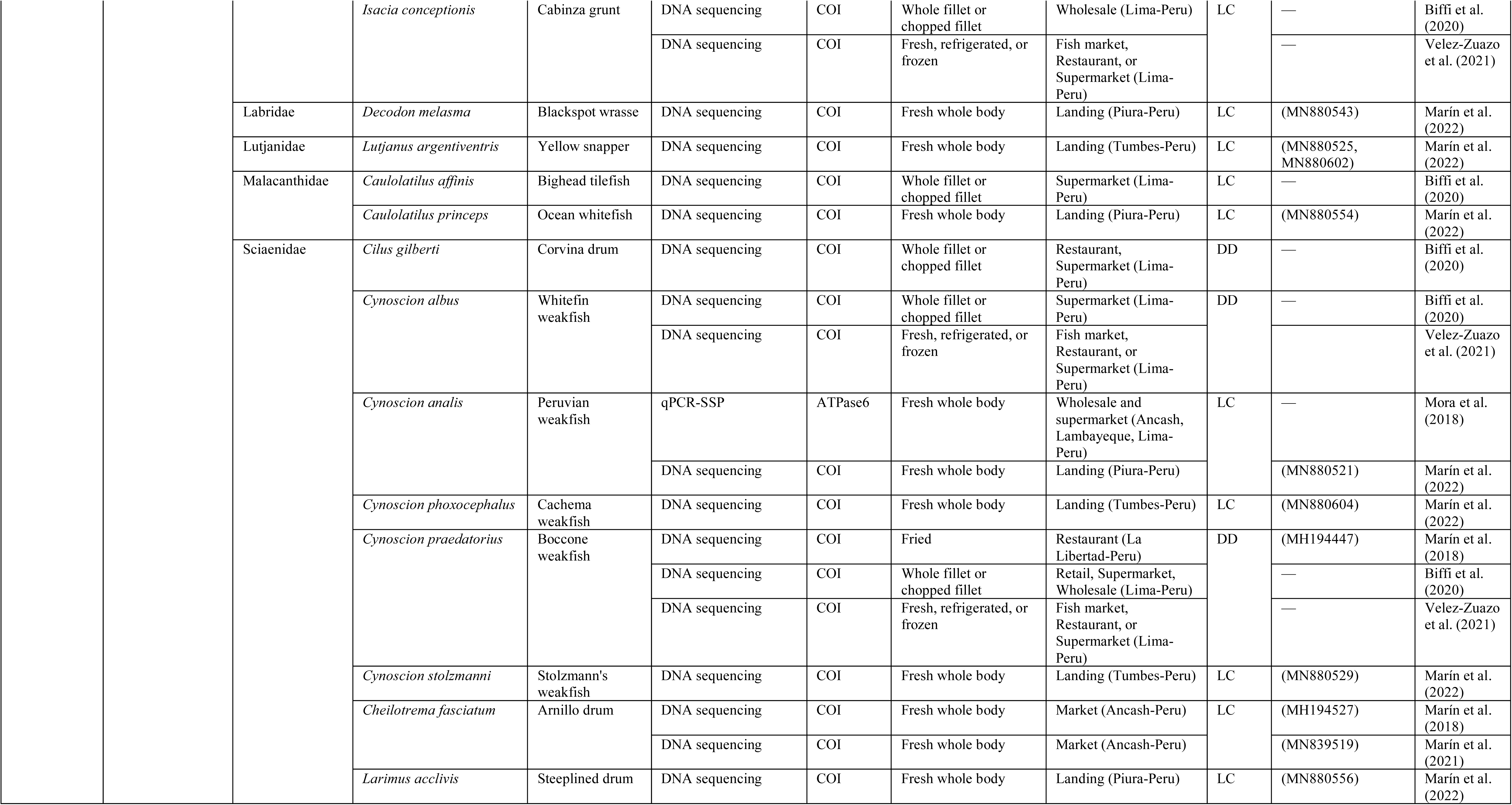
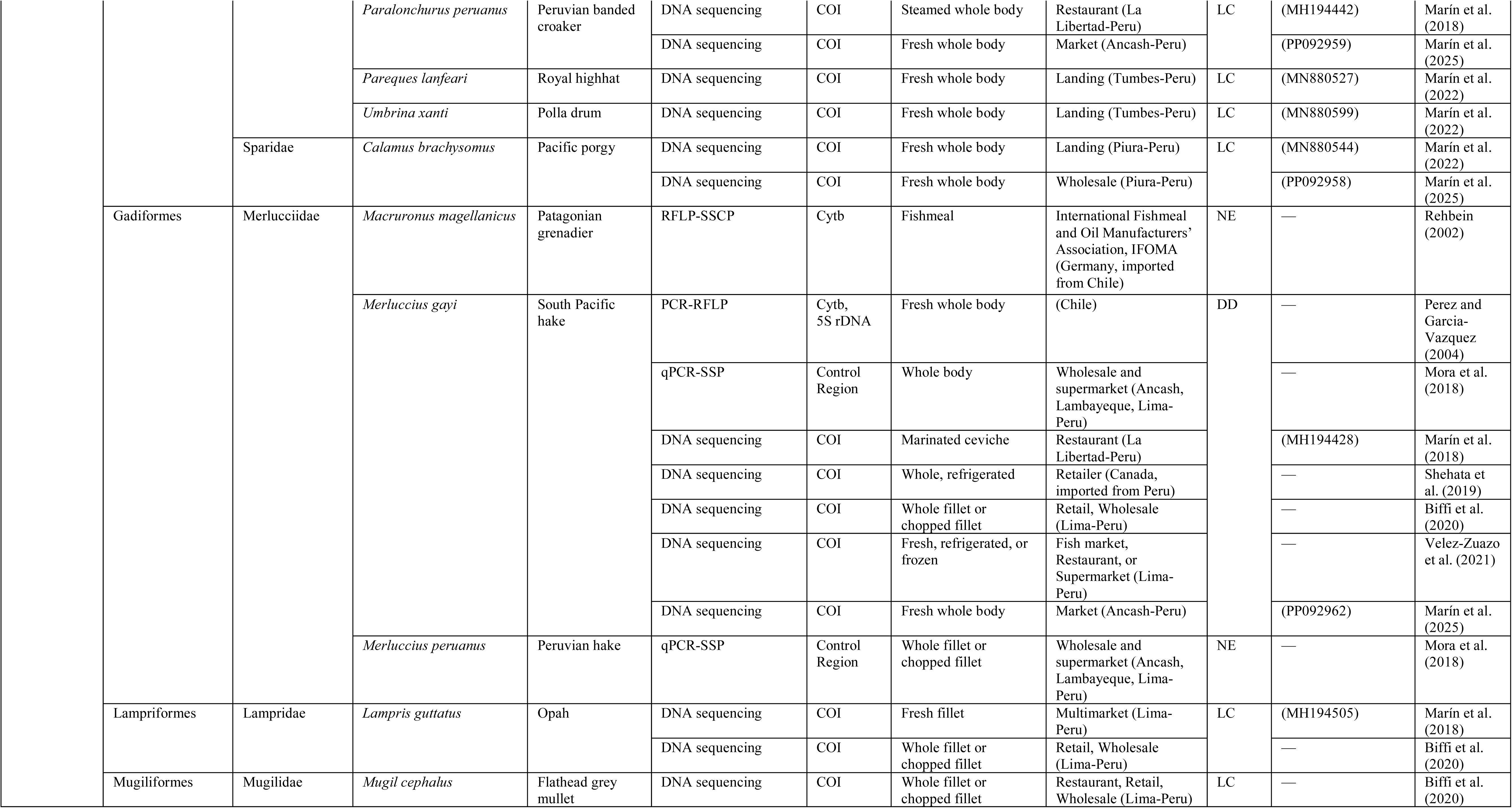
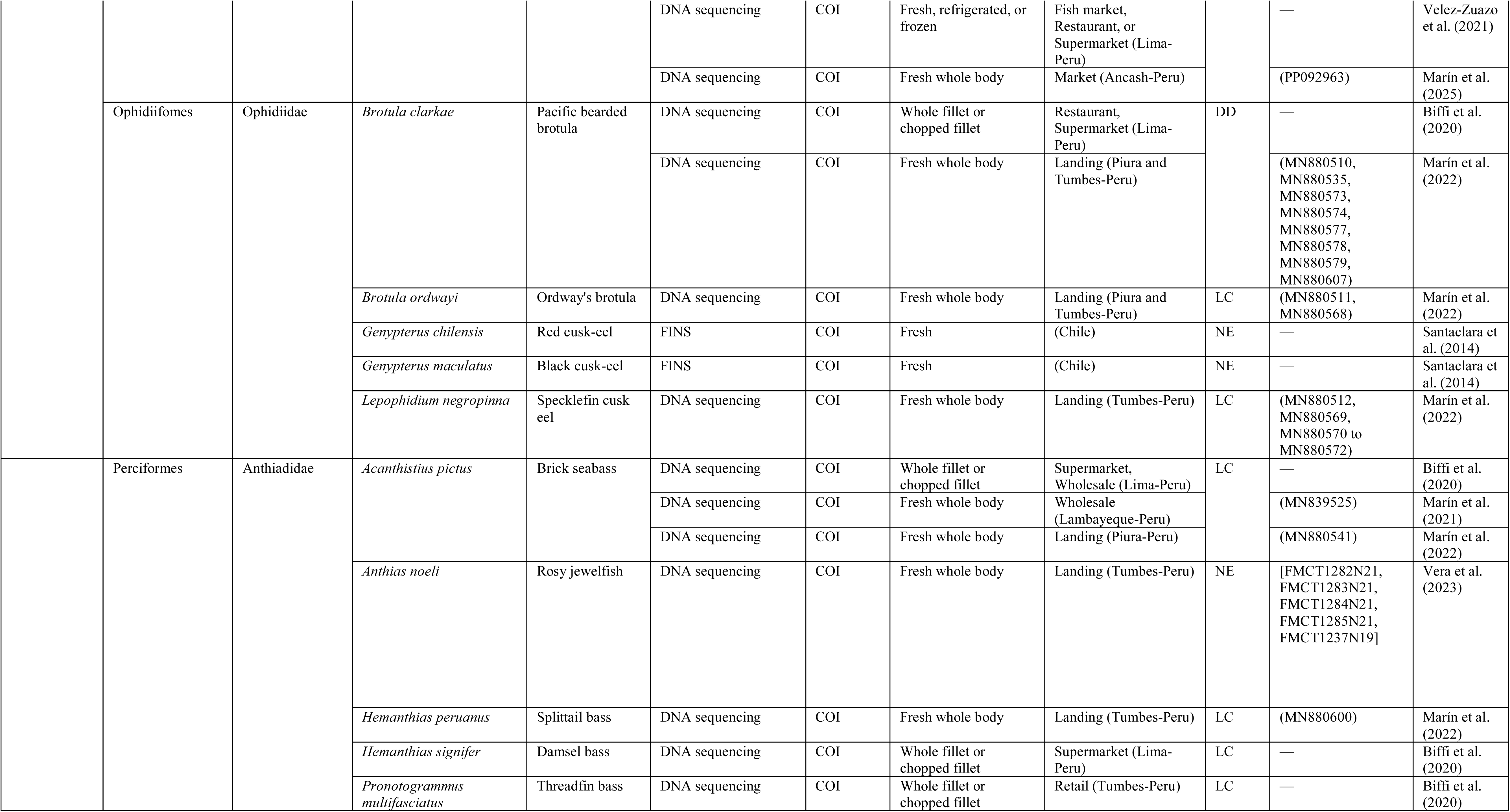
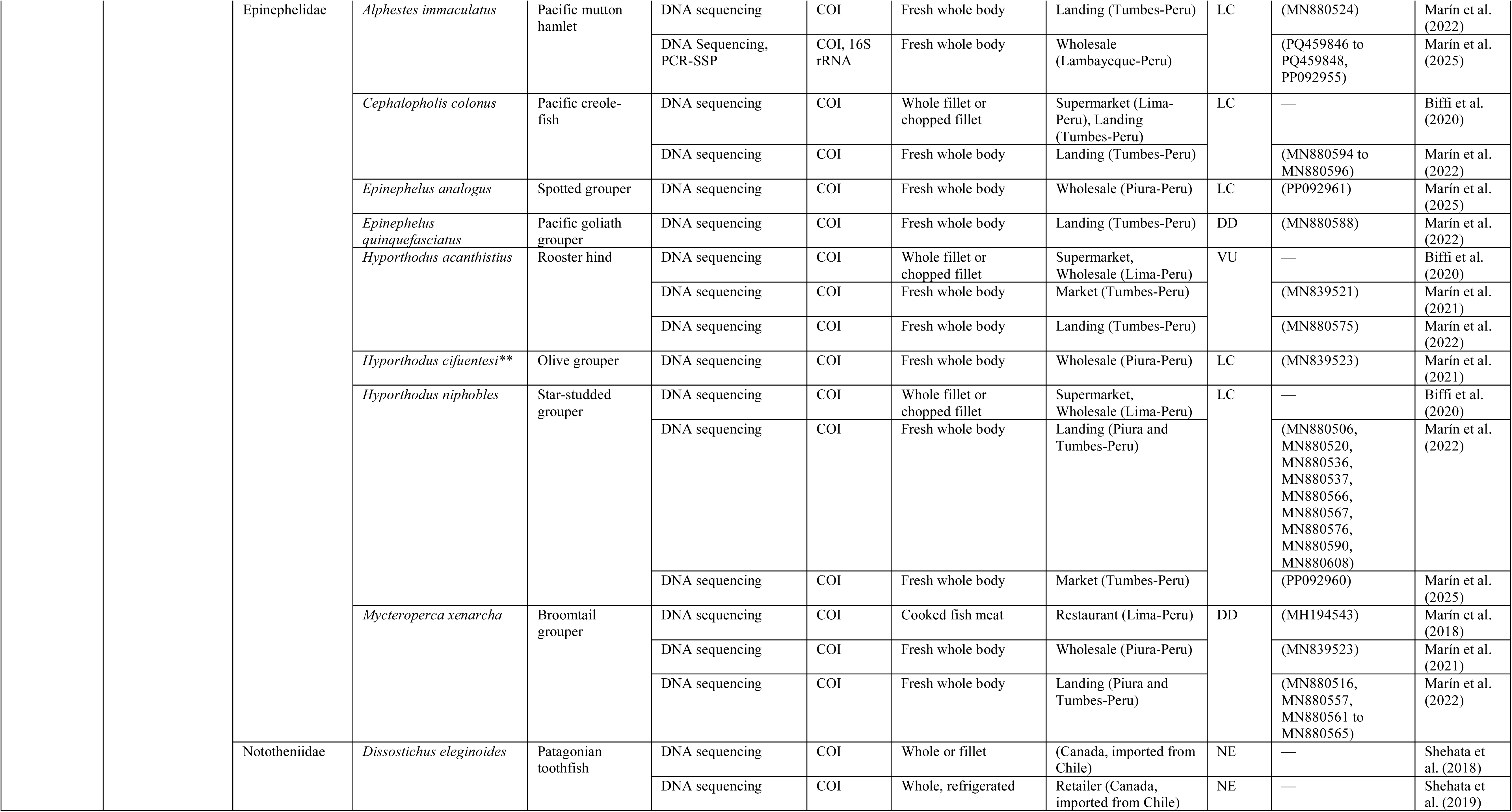
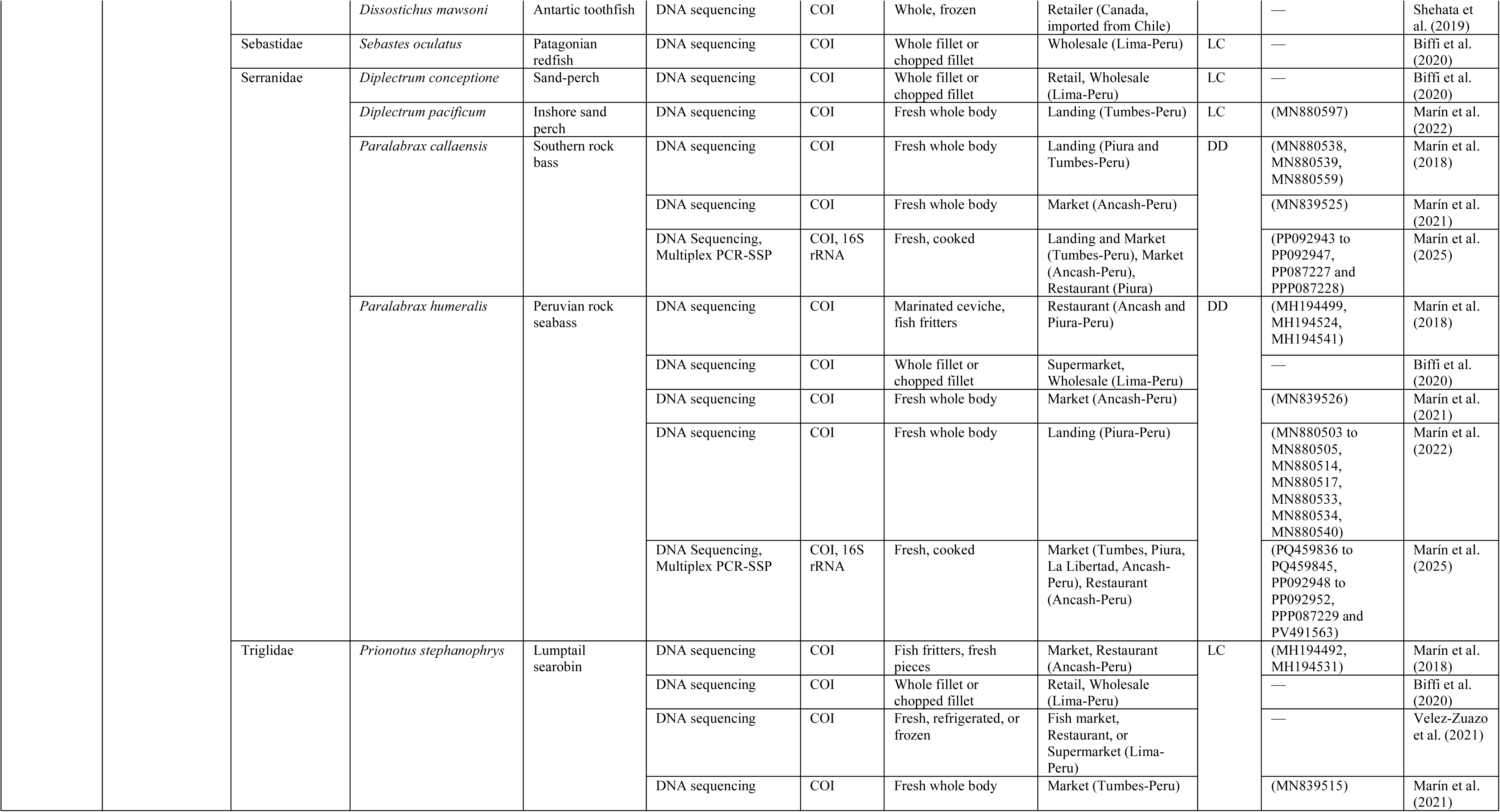
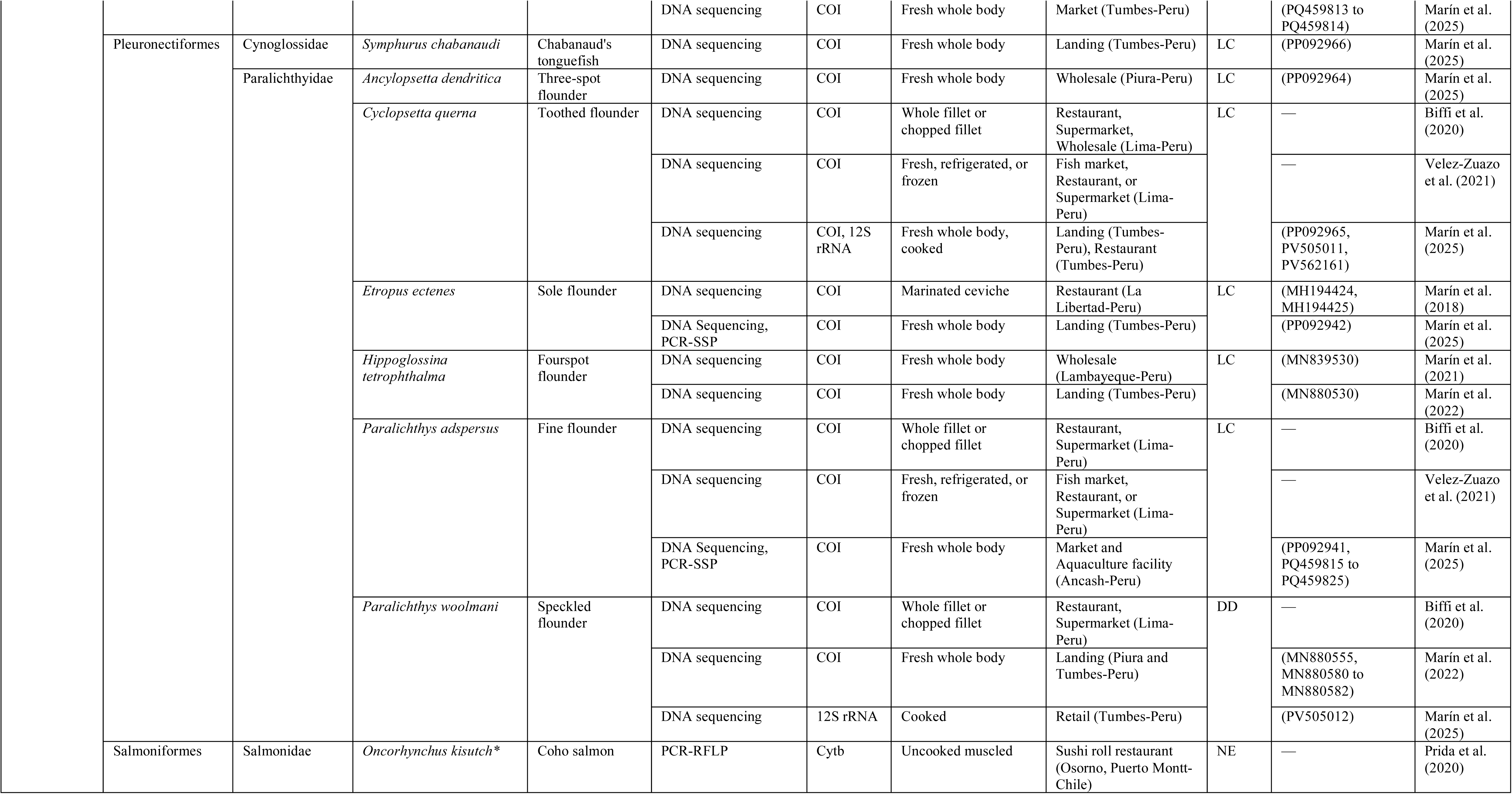
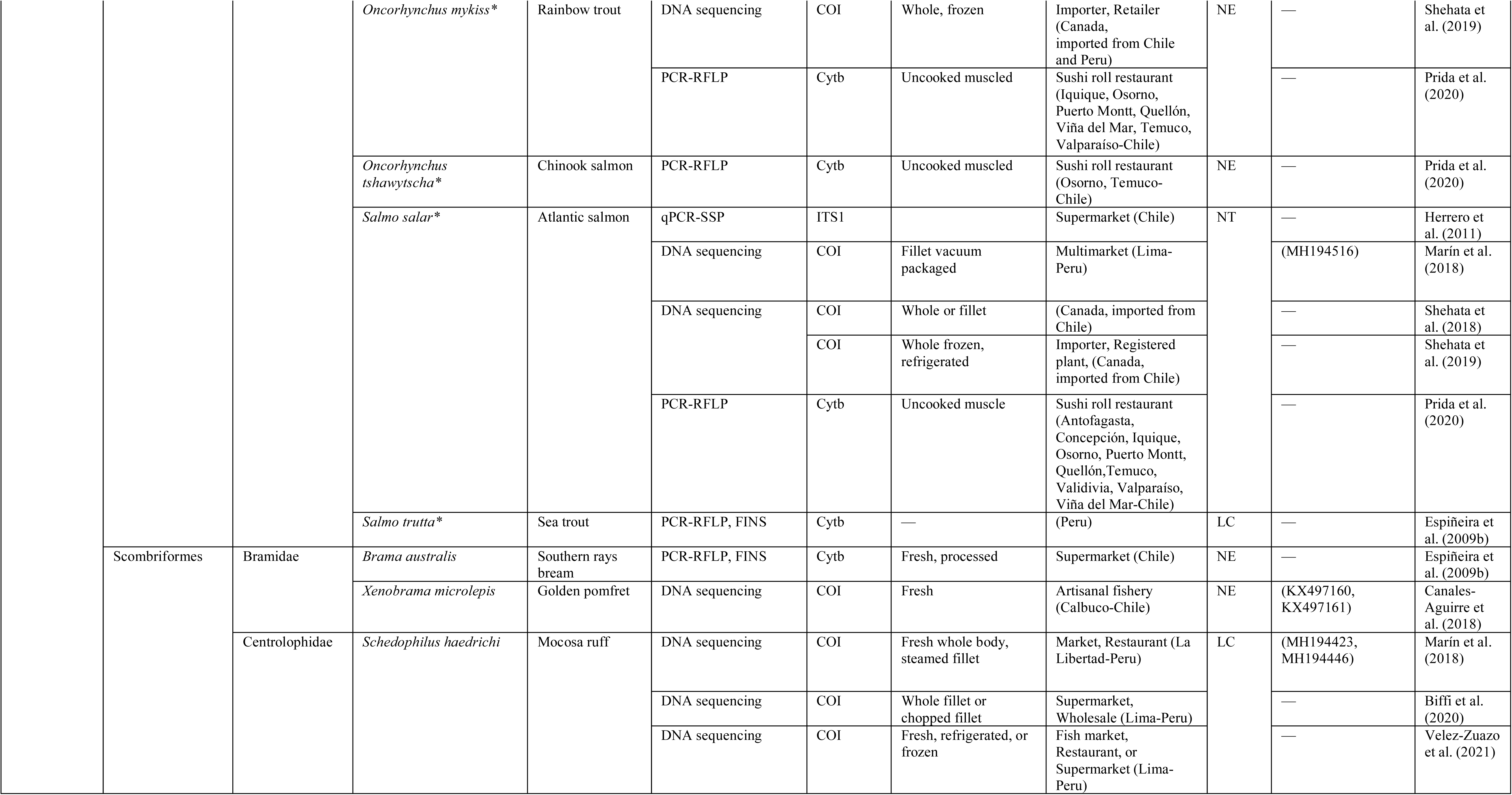
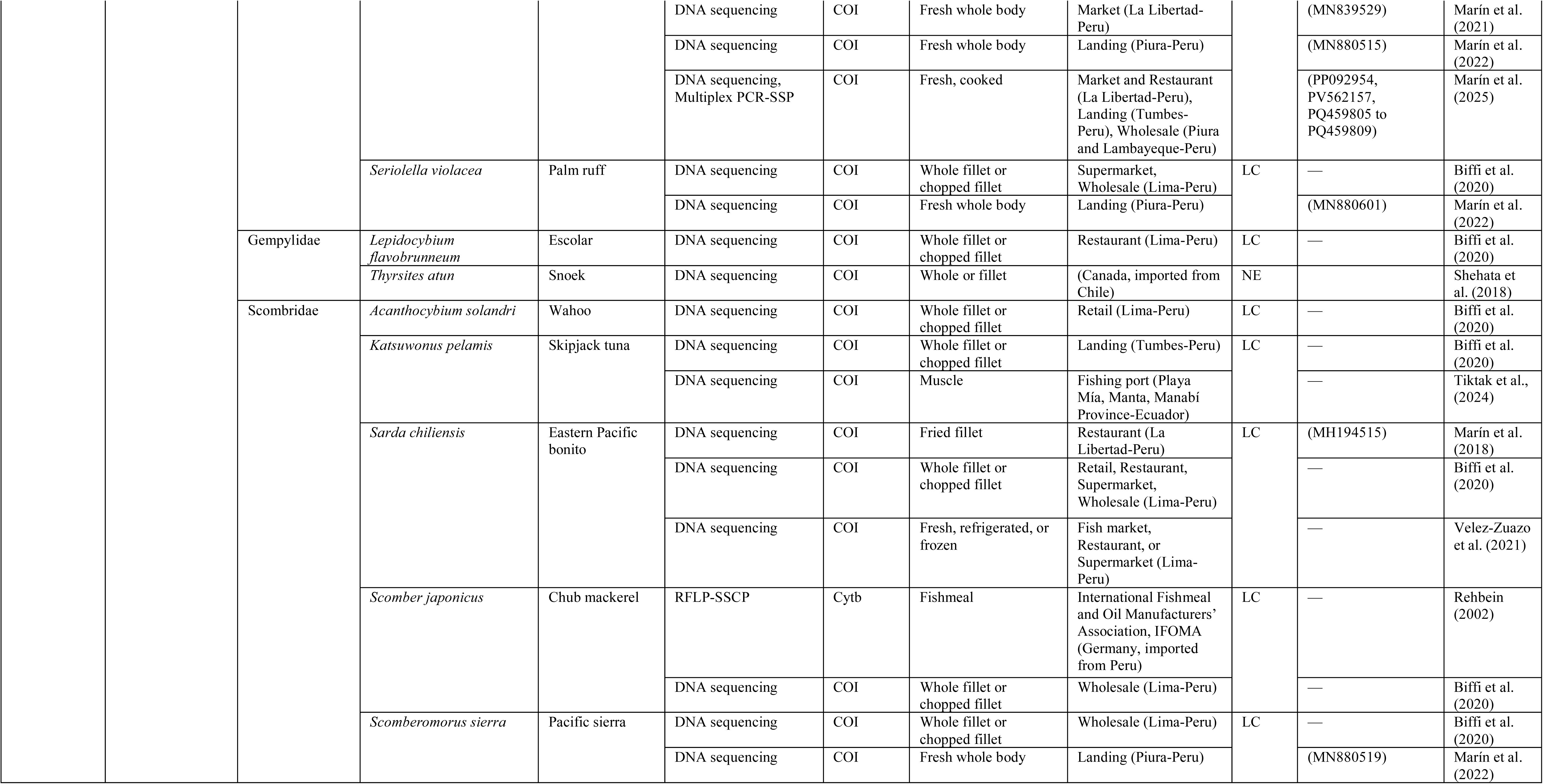
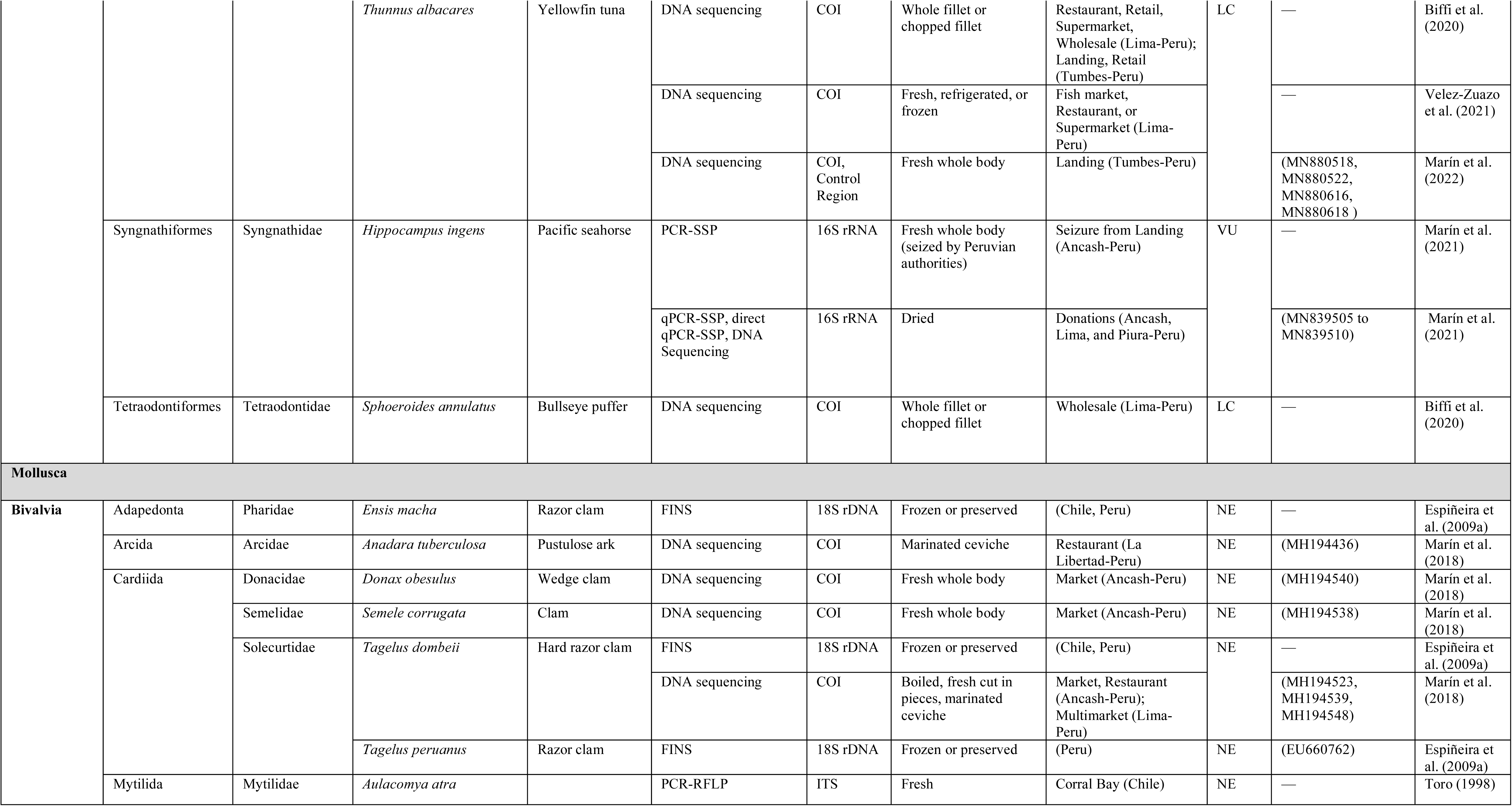
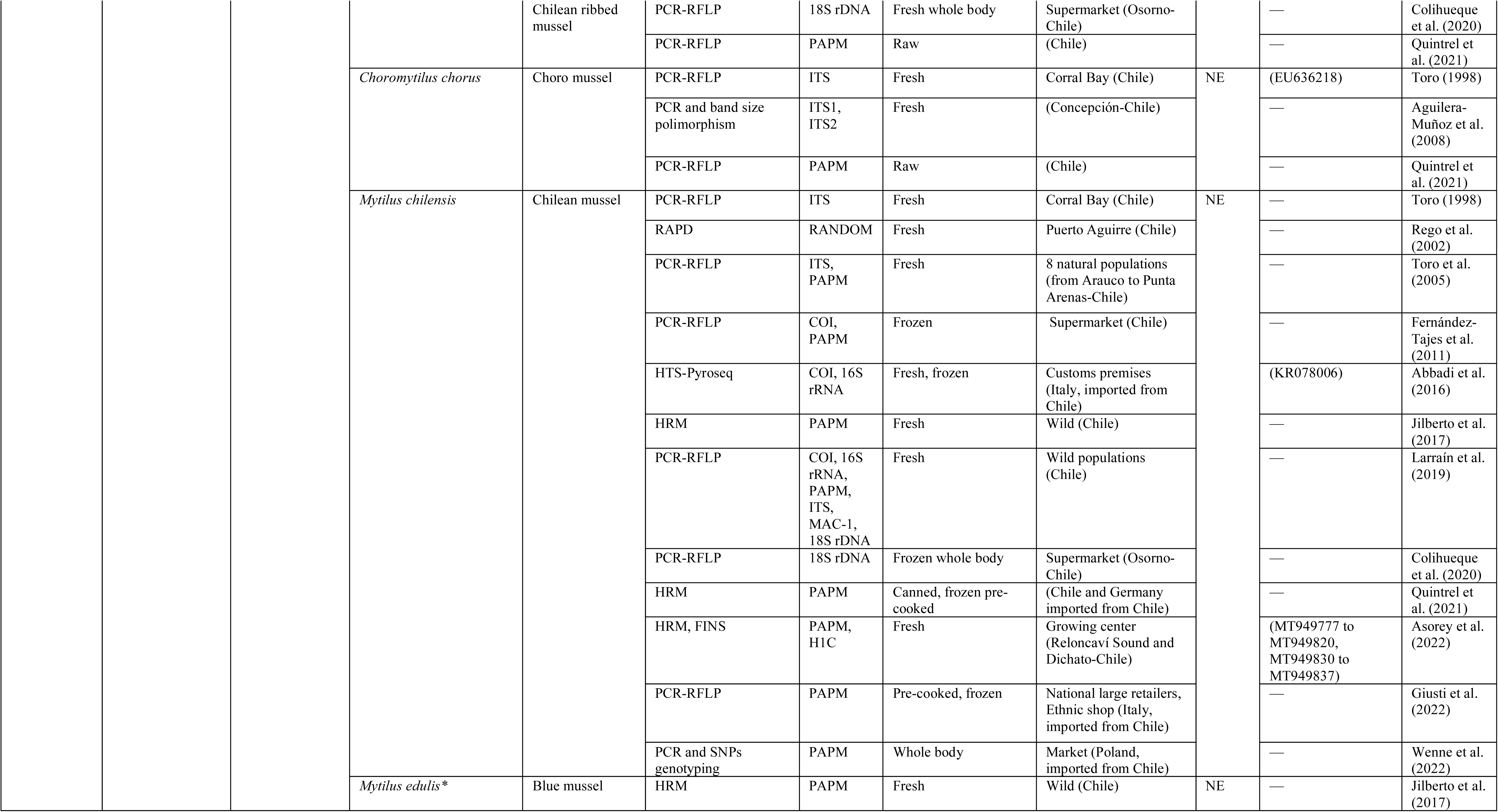
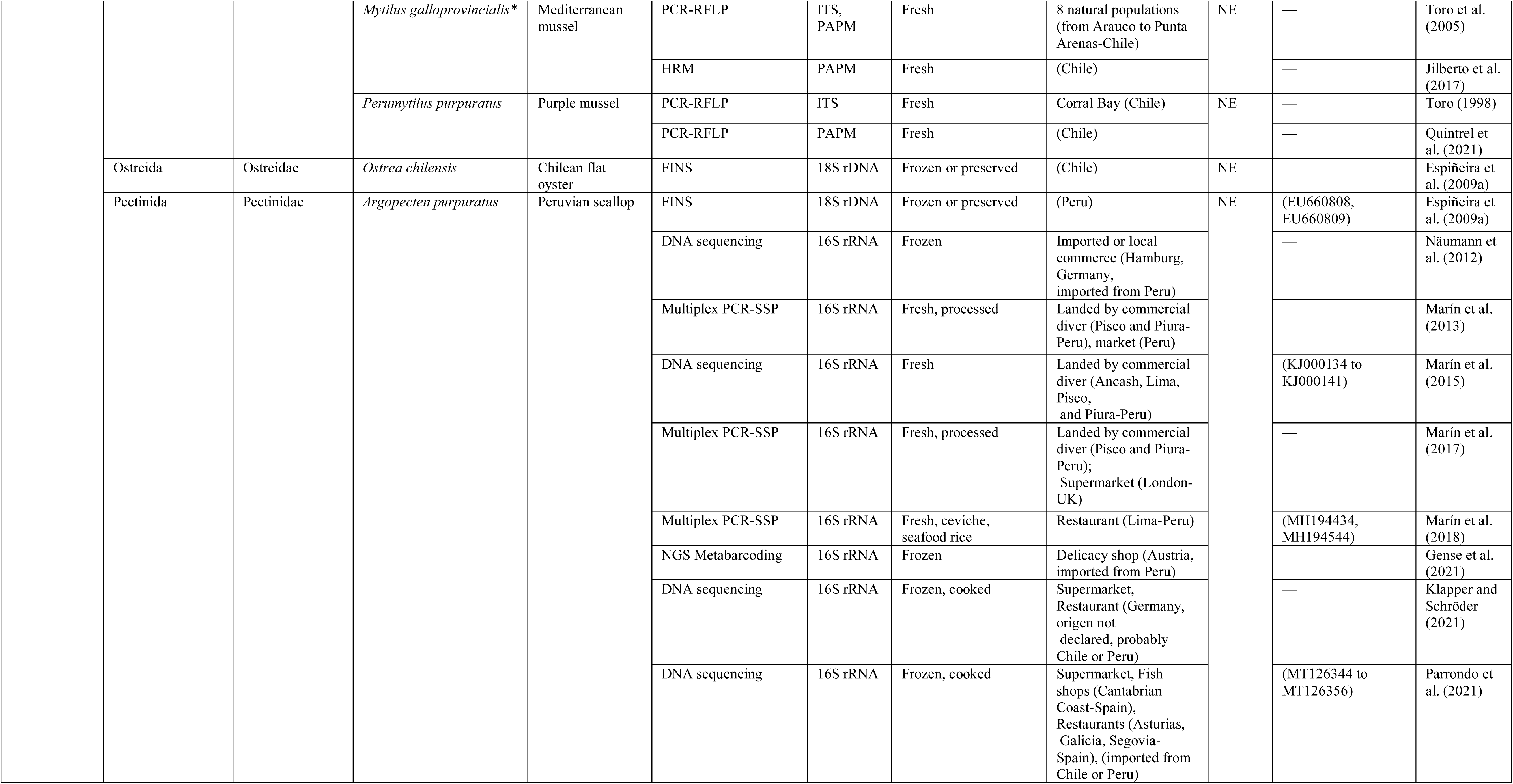
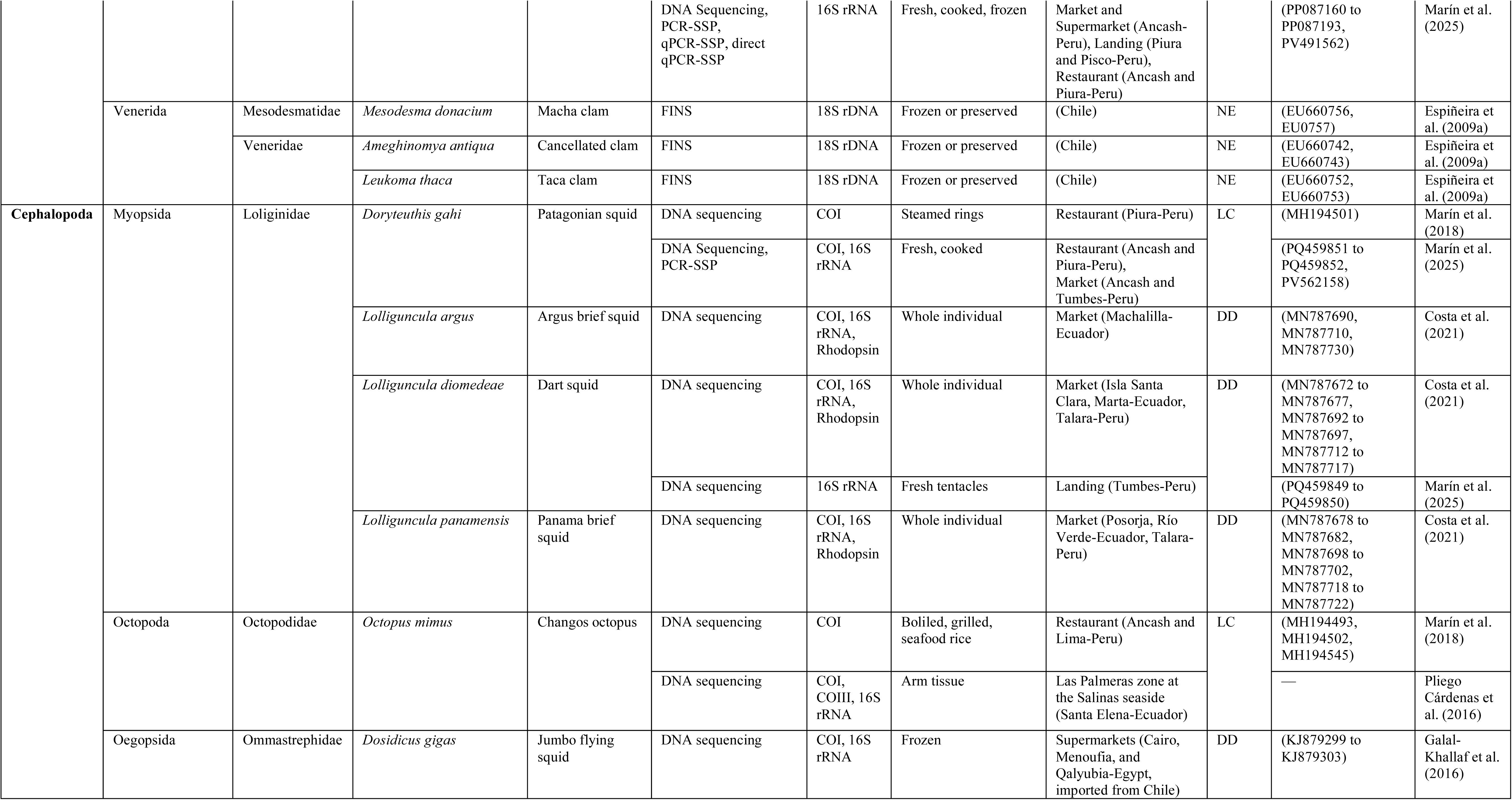
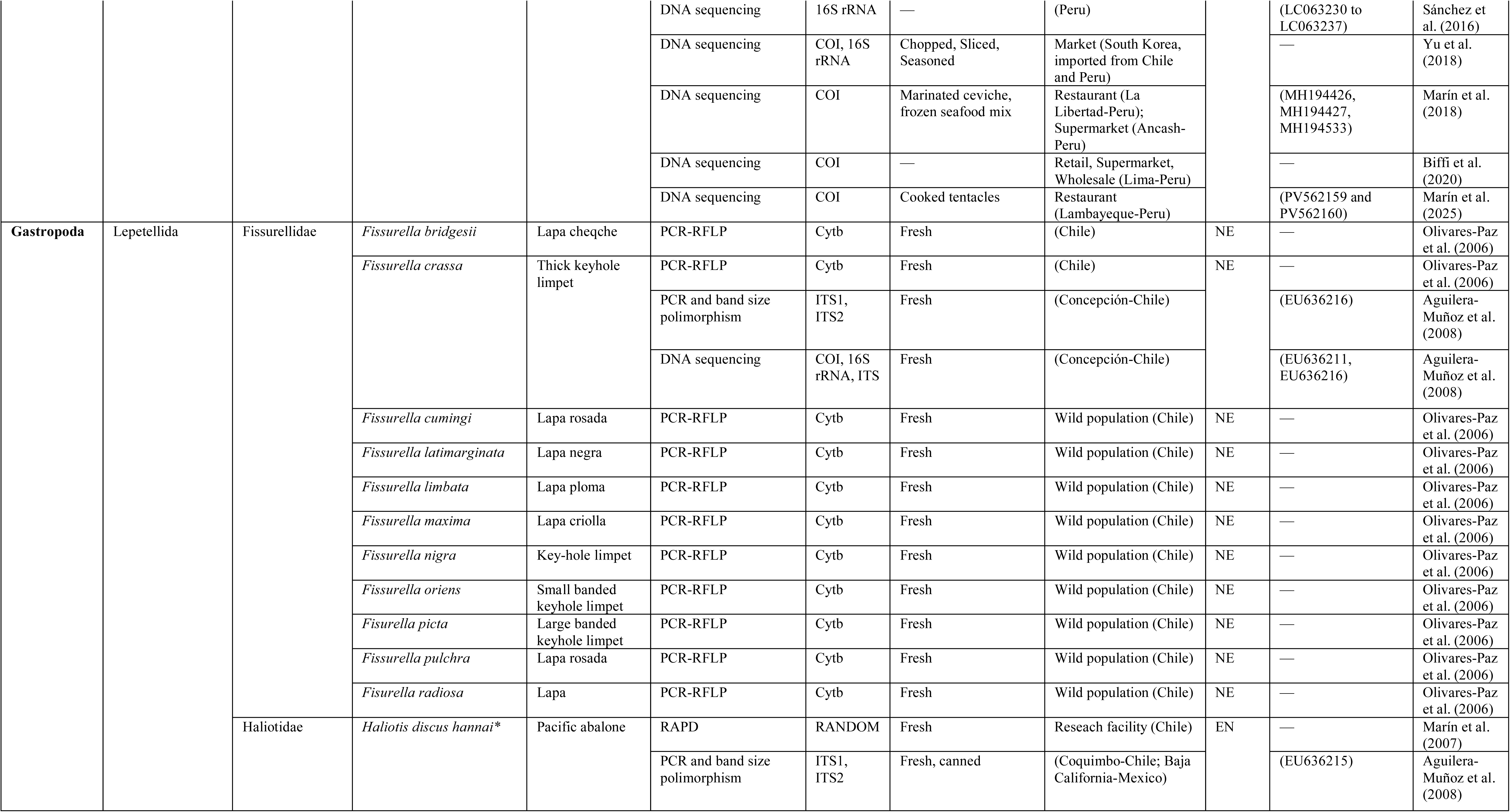
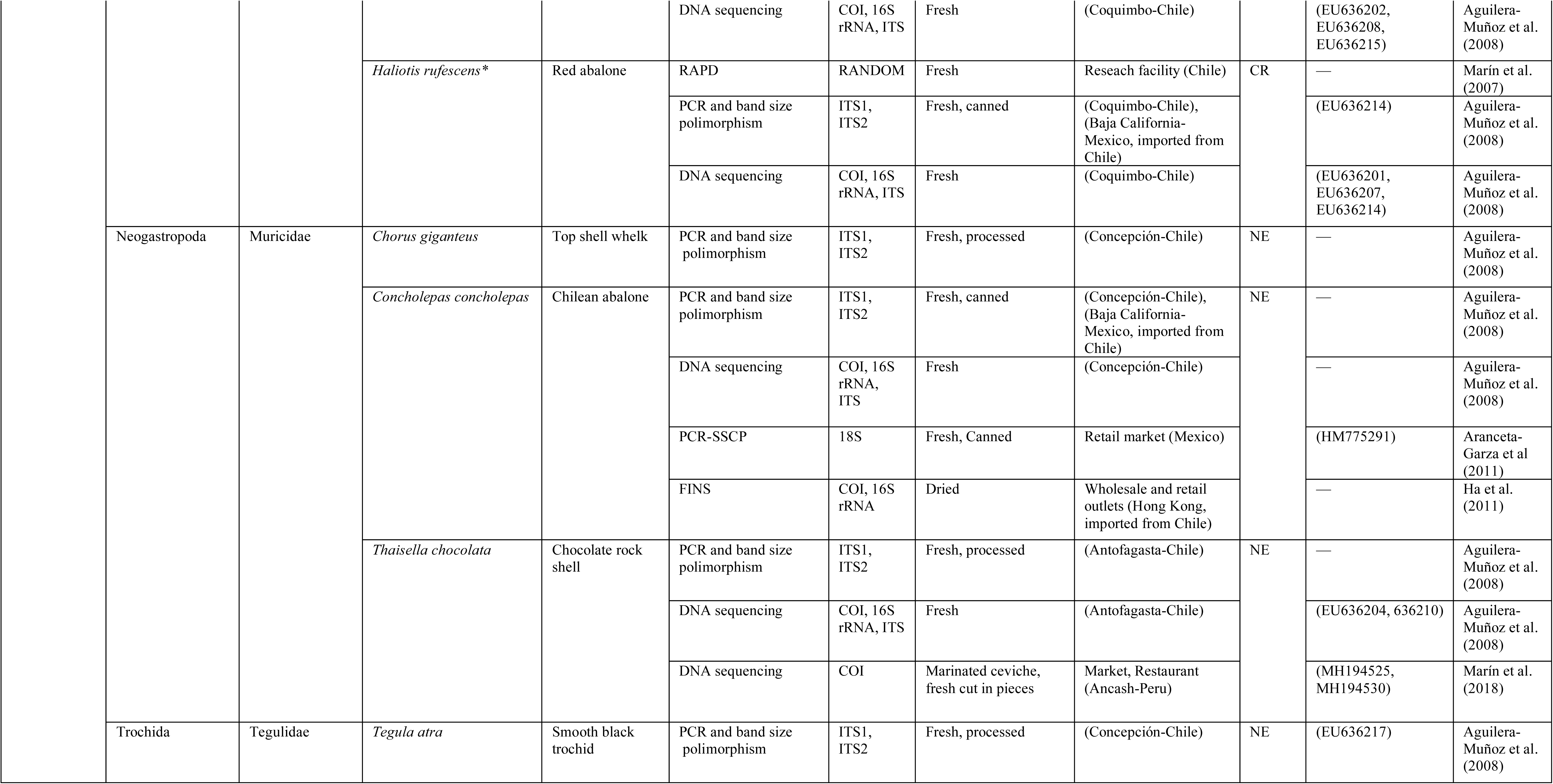
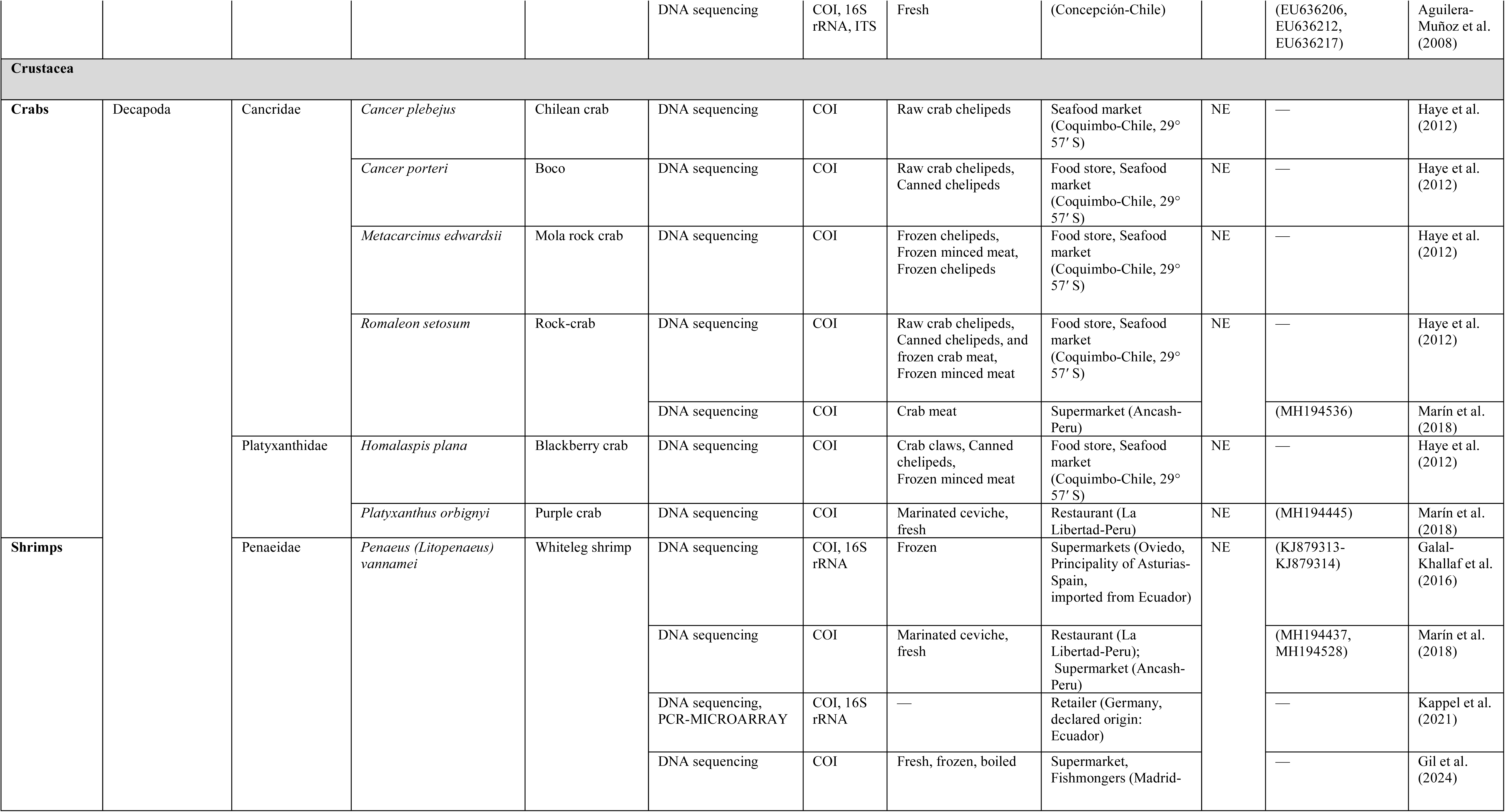
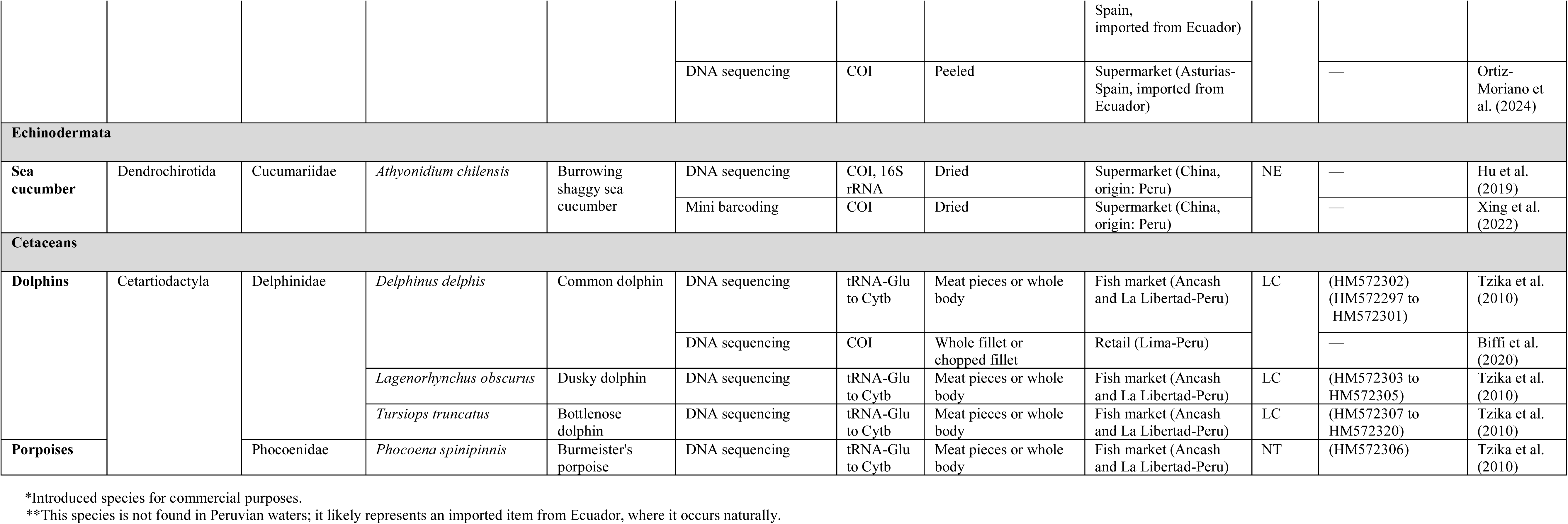
Species diversity found in the articles reviewed by the present study.

#### 3.4.1 Fish

Fish were the most extensively studied seafood group, with samples analyzed in 36 studies. A total of 152 species were identified, distributed within 59 families and 28 orders. This group accounting for 74.5% of all identified species. The diversity within the fish group was primarily dominated by finsfish, which included 104 species (68.4%). This was followed by sharks with 30 species (19.7%), batoids with 17 species (11.2%), and chimaeras with just 1 species (0.7%). The fish group with the highest number of identified species belongs to the shark *Carcharhinus*, which has 7 species. This was followed by the finfish genus *Cynoscion*, which has 5 species.

#### 3.4.2 Mollusks

The mollusk group comprised 40 species identified in 32 studies, distributed across 17 families and 13 orders. This group was evenly represented by gastropods and bivalves, with each group containing 17 species (42.5% each). Cephalopods were represented by 6 species, accounting for 15% of the mollusk group. The commercially important Chilean keyhole limpet species belonging to the genus *Fissurella* accounted for most of the gastropod species (12 out of 17 species).

#### 3.4.3 Crustaceans

This shellfish group was analyzed in seven studies, including seven species from three families within one order. Six crab species, accounting for 85.7% of the findings, and one shrimp species, making up 14.3%, were identified in the studies reviewed herein.

#### 3.4.4 Cetaceans

Cetaceans were represented by only four species (2%) in two studies conducted in Peru. The first study, reported by Tzika et al. (2010), analyzed DNA sequences of the mitochondrial CYTB gene from meat samples purchased in local markets across various coastal cities in Peru. This study identified four cetacean species namely: *Delphinus delphis*, *Lagenorhynchus obscurus*, *Phocoena spinipinnis*, and *Tursiops truncatus*. I should mention that *D. capensis* was also reported but it was later synonymized with *D. delphis* in 2015 (see Cunha et al., 2015). The second article, published by Biffi et al. (2020), provided molecular evidence of dolphin meat sold in a market in Lima, the capital of Peru. Their findings, based on COI sequences, revealed the illegal trade of a *D. delphis* meat sample labeled as “*muchame*”, which is the market term used in Peru for fresh or cured cetacean meat. These results indicate that illegal catches and trade of small cetaceans continue to occur in Peru, despite conservation efforts and a ban against on the capture and commercialization of these species established by the Peruvian government in Law N° 26585 on April 9, 1996 (El Peruano, 1996). This situation aligns with previous findings reported by Campbell et al. (2020).

#### 3.4.5 Holothurians

This animal group included a single species, the burrowing shaggy sea cucumber (*Athyonidium chilensis*), which accounted for the smallest proportion (0.5%) of the total number of species analyzed in this review. Only two studies conducted in China (Hu et al., 2019; Xing et al., 2021) have focused on this commercially valuable group, underscoring the need for more research from ESPO countries to enhance our understanding of local holothurian diversity.

### 3.5 Limitations of DNA-based methods in identifying ESPO species

While most DNA-based methods for identifying seafood species are highly sensitive, efficient, and accurate, several factors can complicate the identification process. These include:

- **Degraded DNA**: highly degraded DNA, often found in canned and cooked seafood due to physical and chemical alterations, can hinder DNA extraction and further PCR amplification. DNA mini-barcoding may help analyze degraded DNA, but it can struggle to distinguish closely related species due to low coverage in gene regions with minimal or no variation. This limitation often restricts identification to the genus level or higher taxonomic categories.
- **Non-target DNA amplification**: Universal primers, which are frequently used for seafood species identification, can mistakenly bind to non-target DNA, including that of parasites and bacteria associated with the target organism.
- **Lack of a comprehensive database with reference sequences**: a database containing low-coverage of reference sequences for species diversity can negatively impact seafood identification. The success of DNA sequencing methods for species identification relies heavily on a comprehensive DNA sequence database for accurate sequence comparison.
- **Incorrect marker selection**: choosing the right DNA marker is crucial for successful seafood species identification. For example, slow-evolving markers may fail to distinguish between closely related or recently diverged species. Additionally, low copy number markers, such as those from the nuclear genome, can complicate the identification of highly processed seafood products or when only a small amount of tissue is available.
- **Poor primer design:** effective primer design is essential for successful PCR amplification, and this requires the use of appropriate bioinformatics tools and experienced researchers. Primers that are designed based on highly variable regions or on a limited number of sequences may inadvertently miss detecting crucial mutations within the template strand that correspond to the complementary region of the primer’s 3’ end. This oversight can inhibit PCR amplification in organisms that contains these mutations.
- **Intrinsic biological nature of target seafood species**: the biological complexity of target seafood species can also complicate identification. Factors such as species complexes, cryptic taxa, recently diverged species, hybridization, and interspecific conservatism can hinder accurate interpretation of seafood species identification results.
- **Human error**: another important factor to consider during the identification process is the human error factor. These errors can occur at any stage, including seafood sampling, molecular identification analysis (such as DNA extraction, PCR amplification, and sequencing), and interpretation of the results.

Reviewed papers have highlighted some challenges in the successful identification of certain seafood species from the ESPO. These challenges primarily stem from a lack of comprehensive reference sequences and low levels of genetic differentiation among closely related species. Reviewed studies reported several instances of identifications at the genus level for tunas (*Thunnus* sp.), smooth-hound sharks (*Mustelus* sp.), rays (*Gymnura* sp. and *Myliobatis* sp.), silversides (*Odontesthes* sp.), and clams (*Semele* sp.) (Marín et al., 2018; Biffi et al., 2020; Velez-Zuazo et al., 2021). Furthermore, species complexes have complicated the identification of butterfly rays (*Gymnura* sp., Ehemann et al., 2024b), octopuses (*Octopus* sp., Marín et al., 2018), and keyhole limpets (*Fissurella* sp., Olivares-Paz et al., 2006). These findings underscore the urgent need for developing an extensive database of DNA sequences for seafood species in the ESPO. To address these issues, researchers should consider exploring alternative and more robust molecular techniques, like multilocus approaches (Feitosa et al., 2018), phylomitogenomics (Alfaro et al., 2025), and whole genome sequencing (WGS) combined with RAD-Seq targeting species-specific SNPs (Jiang et al., 2020).

### 3.6 IUCN conservation status

According to the International Union for Conservation of Naturés Red List of Threatened Species, out of the 204 species analyzed in the studies reviewed, 11 are listed as Critically Endangered (CR) (5.4%), 11 as Endangered (EN) (5.4%), 22 as Vulnerable (VU) (10.8%), 9 as Near Threatened (NT) (4.4%), 80 as Least Concern (LC) (39.2%), 17 as Data Deficient (DD) (8.3%), and 54 as Not Evaluated (NE) (26.5%) (see Fig. 2). Threatened categories (CR, EN, or VU) include a total of 44 species (21.6%), with sharks accounting for the majority at 24 species (54.6%), followed by 13 batoids (29.5%), 4 finfish (9.1%), 2 gastropods (4.5%), and 1 chimaeroid fish (2.3%) (Fig. 2).

These findings highlight the rapid depletion of several commercial elasmobranch species, primarily due to intense fishing pressure, overexploitation, and high bycatch rates. Additionally, the K-strategist nature of these species (low fecundity, long gestation periods, slow growth, and late sexual maturity) makes them particularly vulnerable to even moderate levels of fishing pressure (Wheeler et al., 2020). All these characteristics contribute to their sensitivity to environmental changes and human activities (Marín et al., 2018).

On the other hand, the LC category primarily consists of finfish species, with 71 out of a total of 80 LC species. This category includes various small to medium-sized fish species, such as anchovies, herring, mullets, pilchards, and sardines. These species exhibit r-strategist characteristics, which include high fecundity, early age at first reproduction, and short life spans, enabling them to withstand high levels of fishing pressure (Crosetti, 2016; Patti et al., 2020; Owatari et al., 2024).

Of particular concern is that a considerable proportion of the species compiled are categorized as NE with 54 species (26.5%) falling into this group. This category mainly comprises shellfish species, including 40 species in total: 17 bivalves, 15 gastropods, 7 crustaceans, and 1 holothurian, along with 13 fish species. These findings indicate an urgent need for population and demographic studies of commercially important shellfish species from the ESPO region. The results from such studies would contribute to better assessments and determinations of the current conservation statuses and population trends, ultimately leading to improved management and sustainable exploitation of those resources.

### 3.7 Mislabeling meta-analysis in ESPO seafood species

This study presents the first quantitative meta-analysis focused on seafood mislabeling of ESPO species sold globally. Based on the Level 3 screening criteria, 39 articles were selected, which included a total of 1,806 cured samples purchased throughout the entire supply chain, from harvesting and processing to distribution and retail, across 16 countries. The number of samples for each taxonomic group were: 882 finfish (48.8%), 485 bivalves (26.9%), 337 sharks (18.7%), 39 cephalopods (2.2%), 26 batoids (1.4%), 27 crustaceans (1.5%), 6 gastropods (0.3%), 2 holothurians (0.1%), 1 chimaera fish (0.05%), and 1 cetacean (0.05%).

A re-analysis of seafood samples with publicly available DNA sequences allowed me to upgrade eight seafood samples that were previously identified only to the genus level due to a lack of reference sequences at the time of their publication. Among these upgrades was a canned sardine identified by Jérôme et al. (2003b), which was reclassified from *Sardinops* sp. to *S. sagax* (GenBank AY394046). Furthermore, four samples originally identified as *Mustelus* sp. were upgraded to *Mustelus whitneyi* (Genbank MH194477, MH194478, MH194497, and MH194506). Additionally, two samples identified as *Gymnura* sp. were upgraded to *Gymnura afuerae* (Genbank MH194463 and MH194465), and one sample initially identified as *Semele* sp. was reclassified as *Semele corrugata* (Genbank MH194538) (Marín et al., 2018). This findings highlight the importance of sharing DNA records, which aligns with the data availability policies of several relevant journals. Sharing DNA records not only supports the conclusions of studies but also benefit the research community by allowing re-analysis of published results, facilitating meta-analyses, and reducing costs of further research (Sielemann et al., 2020). In this review, it was noted that out of the 39 articles selected for the meta-analysis, 27 studies focused on DNA sequencing, of which only 11 made their DNA records publicly accessible.

The results of the global mislabeling analysis are shown in Fig. 3. These results indicated that out of the 1,806 seafood products, 448 were mislabeled. This represents a global mislabeling rate of 24.8% (95% CI [22.9-26.9]). The mislabeling rate found in this review, which focused specifically on ESPO seafood species, falls within the range of previous global mislabeling meta-analyses, which reported rates between 19% and 30% (Pardo et al., 2016; Warner et al., 2016; Luque & Donlan, 2019). The complete meta-dataset can be found in Supplementary Table S1. While the findings of the mislabeling meta-analysis are informative, they should be interpreted with caution due to varying sampling strategies and efforts used across the different studies reviewed.

**Fig. 3.**
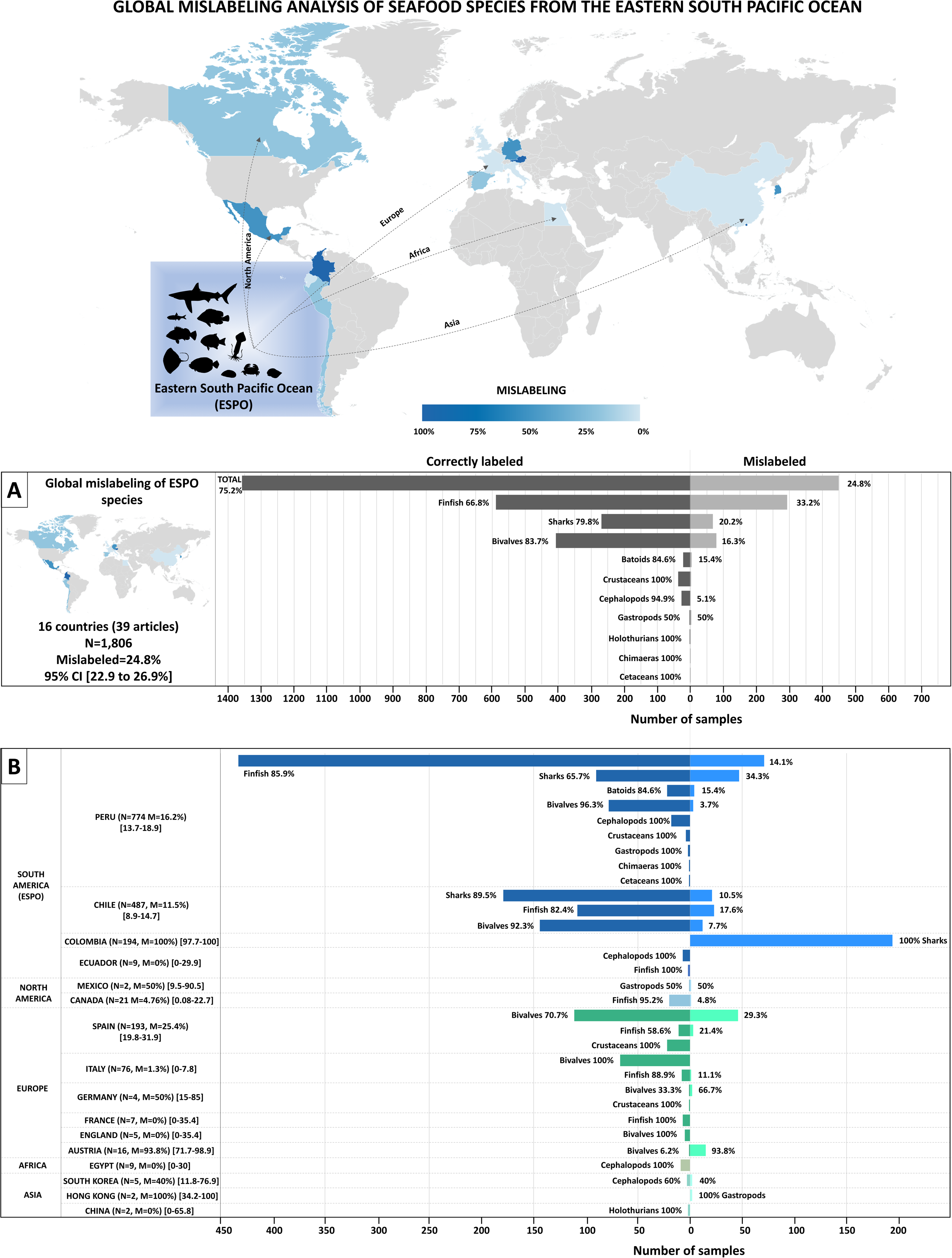
Results of the global meta-analysis for seafood from the Eastern South Pacific Ocean (ESPO). The heat map illustrates the countries from which seafood samples were collected and analyzed. Color intensity indicates the mislabeling rate calculated for each country, as displayed in the mislabeling rate color scale. The back-to-back bar graph in Panel A presents the overall global rates of mislabeled and correctly labeled samples, along with rates specific to each seafood category. The back-to-back bar graph in Panel B depicts the proportions of mislabeled and correctly labeled samples observed in individual countries, categorized by seafood type. Confidence intervals (95% CI) for the mislabeling rates in each country are provided in brackets. “N” indicates sample size, and “M” denotes the mislabeling rate.

#### 3.7.1 Mislabeling per geographic region

Overall, the reviewed samples were collected from five continents, encompassing 16 countries. The mislabeling results, categorized by continent, indicated that Asia had the highest rate of mislabeling at 44.4% (4/9, CI [18.9-73.3]), followed by South America at 25.6% (375/1464, 95% CI [23.7-28.2]), Europe at 22.3%, (67/301, 95% CI [17.9-27.3]), and North America at 8.7%, (2/23, 95% CI [2.4-26.8]).

No mislabeling was detected in samples from Africa (0/9, 95% CI [0-30]), although the sample size for this continent was small and not representative (see panel B in Supplementary Table S1). Only South America and Europe had sampling sizes exceeding the threshold of n > 25, consequently only both continents were included in the statistical test of significant association. The mislabeling rates observed in these two continents did not show a statistically significant relationship (Chi-square, *p* = 0.25) (Fig. 4).

**Fig. 4.**
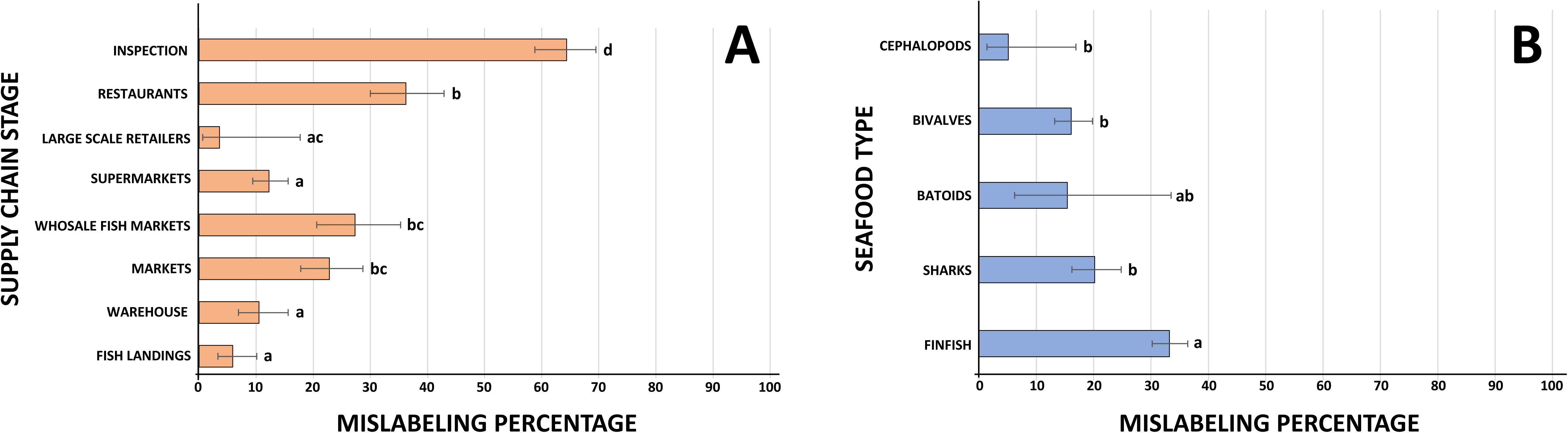
Mislabeling rates by A) continent and B) country. Bars represent the mislabeling rate, error bars indicate the 95% confidence intervals. The relationship between the mislabeling rates in Europe and South America showed no significant deviation. Bars labeled with different lowercase letters are significantly different at *p* < 0.05. Statistical significance was determined using either the Chi-square test or Fisher’s exact test, depending on the sample size.

When mislabeling analyses were organized by country, the highest mislabeling rates of 100% were found in Colombia (194/194, CI [97.7-100]) and Hong Kong (2/2, 95% CI [34.2-100]). It should be noted that samples analyzed from Colombia involved 194 shark fins that were incorrectly labeled as fish bladders. All of these selected fins were confiscated during an official inspection at an airport, therefore mislabeling rates in these samples were expected to be high. On the other hand, the sample size from Hong Kong consisted of only two ESPO samples, which is not representative. Other estimates from countries with small sample sizes (n < 10) reported mislabeling rates of 0% for China, Ecuador, Egypt, England, and France, while Mexico, Germany, and South Korea reported mislabeling between 40% and 50%.

Countries with more representative sample sizes include Peru with a mislabeling rate at 16.5% (125/774, 95% CI [13.7-18.9.3]), Chile at 11.5% (56/487, 95% CI [8.9-14.7]), Spain at 25.4% (49/193, 95% CI [19.8-31.9]), and Italy at 1.3% (1/76, 95% CI [0-7.8]). A statistical test of significant association between the mislabeling rates detected in five countries (Chile, Colombia, Italy, Spain, and Peru) showed non-significant similar relationship only between Chile and Peru (Chi-square, *p* = 0.27). All other relationships showed signs of significant deviation (Fig. 4). This finding was mainly driven by the strong contrast in mislabeling rates observed across the countries analyzed. Colombia exhibited an extremely high misleading rate, while Spain, Peru, and Chile had moderate to high levels, and Italy had a very low mislabeling rate. Detailed mislabeling results for each country are shown in panel B of Supplementary Table S1.

#### 3.7.2 Mislabeling per distribution channel

The analysis revealed that mislabeling rates varied significantly among the different distribution channels. The highest overall mislabeling rate was found in the inspection category at 64% (195/303, 95% CI [58.8-69.5]). This was followed by restaurants at 36% (76/210, 95% CI [30-42.9]), wholesale markets at 27.3% (38/139, 95% CI [20.6-35.3]), markets at 22.8% (52/228, 95% CI [17.8-28.7]), supermarkets at 12.2% (53/434, 95% CI [9.4-15.6]), warehouse at 10.5% (21/200, 95% CI [6.9-15.6]), fish landings at 5.9% (12/203, 95% CI [3.3-10.1]), and large scale retailers at 3.57% (1/28, CI [0.63-17.7]) (Fig. 5). The “market” category encompassed samples collected from the following subsectors: ethnic shops, delicacy shops, fishmongers, open air markets, retail markets, handicraft markets, fish markets, retailers, and local commerce.

**Fig. 5.**
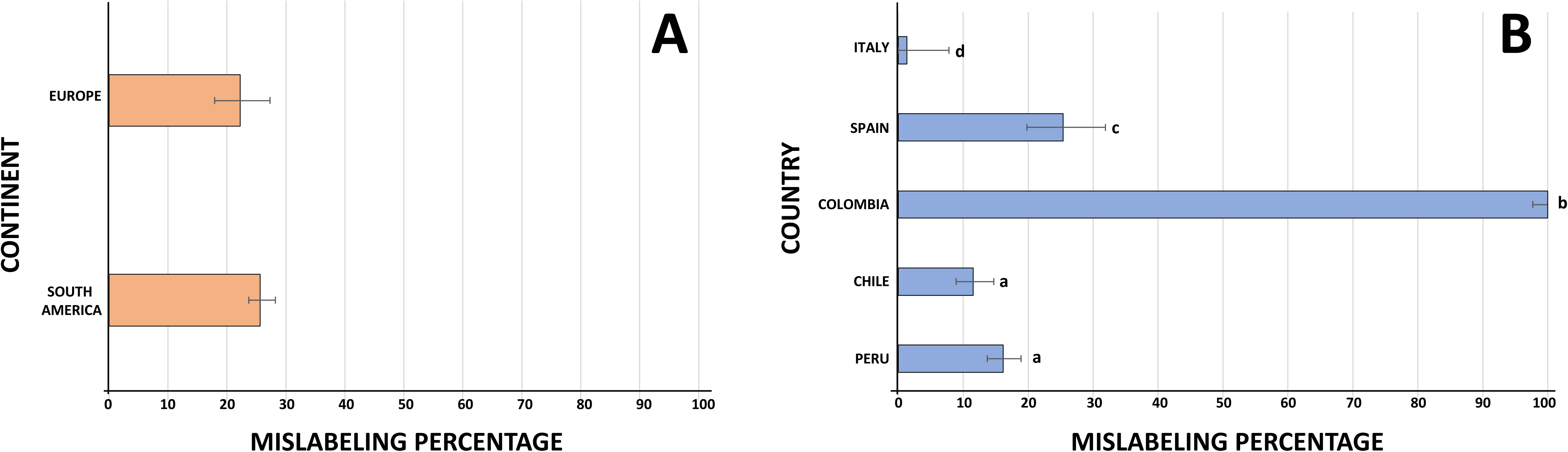
Mislabeling rates by A) supply chain stage and B) seafood type. Bars represent the mislabeling rate, error bars indicate the 95% confidence intervals. Bars labeled with different lowercase letters are significantly different at *p* < 0.05. Statistical significance was determined using either the Chi-square test or Fisher’s exact test, depending on the sample size.

The mislabeling rate observed at the inspection category was significantly higher than in all other categories (Chi-square, *p* < 0.05). This discrepancy can be attributed to the elevated mislabeling rate within this category compared to others. It is important to note that most samples from this category were obtained through seizures at official control points, which tend to contain a higher percentage of fraudulent items. Out of the 303 samples analyzed in this category, 194 (64%) were processed shark fins that were falsely declared as fish bladders at an airport in Colombia (Cardeñosa et al., 2023).

The mislabeling rate at restaurants was significantly higher (Chi-square, *p* < 0.05) compared to other categories, except for wholesale and markets, where the rates were statistically similar (*p* > 0.05). Excluding the inspection category, restaurants exhibited the highest mislabeling rate, while the lowest rate was found at fish landings. This outcome was anticipated, as the initial stages of the supply chain have fewer opportunities for mislabeling. This result aligns with several studies that have shown that mislabeling tends to increase further along the supply chain (Pardo et al., 2016; Marín et al., 2018).

#### 3.7.3 Mislabeling per seafood type

The estimated overall mislabeling rates for the different types of seafood analyzed were highest for gastropods at 50% (3/6, 95% CI [18.8-81.2]), followed by finfish at 33.2% (293/882, 95% CI [30.2-36.4]), sharks at 20.2% (68/337, 95% CI [16.2-24.8], bivalves at 16.1% (78/485, 95% CI [13.2-19.8]), batoids at 15.4% (4/26, 95% CI [6.2-33.5]), and cephalopods at 5.1% (2/39, 95% CI [1.4-16.9]) (Fig. 5). A global Chi-square test indicated a strong significant difference (*p* < 0.05) in the mislabeling rates among the assessed groups (batoids, bivalves, cepahlopods, finfish, and sharks). Subsequent multiple comparison analyses identified significant differences among three groups: finfish vs. cephalopods (Fisher’s exact test, *p* < 0.05), finfish vs. bivalves (Chi-square, *p* < 0.05), and finfish vs. sharks (Chi-square, *p* < 0.05) (Fig. 5). The finfish group exhibited the highest significant mislabeling rate. This is likely due to its high species diversity, the variety of commercial forms, and economic incentives, all of which make this category more susceptible to mislabeling or substitution. In contrast, the smaller sample sizes in other categories, such as batoids and cephalopods, reduce the statistical power to detect meaningful differences. Additionally, the unequal group sizes may influence the robustness of comparisons across seafood categories.

The most frequently encountered substitute species were three threatened shark species. A total of 161 samples, initially labeled as fish bladders (typically sourced from certain finfish species) were actually shark fins from two species: *Carcharhinus falciformis* (n=114, IUCN status: VU) and *Sphyrna lewini* (n=47, IUCN status: CR). The third species, *Prionace glauca* (n=38, IUCN status: NT), was found being used not only as a substitute for other sharks (i.e. hammerheads, shortfin makos, and smooth-hounds) but also for various finfish products (i.e. dolphinfish, rock seabass, and fish bladder). These findings highlight serious conservation concerns, particularly given the involvement of highly threatened species, demanding urgent policy actions and enforcement measurements.

The current review identified a total of 27 Peruvian seafood items labeled as “*tollo de leche*”. Out of these samples, two were finfish, while the remaining 25 belonged to five different shark species. Notably, three of these shark species (*A. pelagicus*, *I. oxyrinchus*, and *S. zygaena*) are classified as threatened by the IUCN. In Peru, the term “*tollo de leche*” was originally designated for a single species: the Critically Endangered humpback smooth-hound shark, *M. whitneyi* (Elliot et al., 1996; Chirichigno & Cornejo, 2001). However, a recent study revealed that this term is now being used to refer to small-sized specimens of various shark species (Núñez & Gozzer-Wuest, 2019). The use of conflicting market names, which constitutes a form of mislabeling (Ahles et al., 2025), can result in overfishing of vulnerable species, hinder the enforcement of regulations, and limit consumers’ ability to make informed choices (Almerón-Souza et al., 2018). To address these issues, efforts should focus on improving labeling accuracy and eliminating conflicting names by creating a standardized list of unique market names (Marín et al., 2018).

Among the invertebrate seafood samples analyzed, the most commonly mislabeled species was the Chilean blue mussel *M. chilensis*. This bivalve was substituted with three different species: *Aulacomya atra*, which was detected in 12 samples collected from a supermarket in Chile (Colihueque et al., 2020), and *M. galloprovincialis* or *M. edulis*, which were found in 15 products from European commerce (Gense et al., 2021). Another bivalve, the Peruvian scallop (*A. purpuratus*), was the most commonly used substitute species. This scallop species was used to replace two types of European scallops: the variegated scallop (*Mimachlamys varia*) in Spain, with 45 substituted samples, and two samples of king scallops labeled as *Pecten maximus* and *Pecten* sp. in Germany. The year-round availability of farmed Peruvian scallop positions this species as a potential substitute for wild, seasonally available local European scallops (Parrondo et al., 2021).

## 4. Conclusions and future perspectives

This research presents the first comprehensive global survey of DNA-based methods used to identify seafood species from the ESPO, with particular focus on species diversity, conservation status, and issues related to mislabeling. The findings from this review contribute significantly to our understanding of past, present, and future trends in molecular methods that support traceability, regulation, and effective management of ESPO species. Among the identification methods, Sanger sequencing technology stands out as the most frequently utilized due to its high efficiency and accuracy. However, faster and more cost-effective techniques, such as species-specific primers and multiplex PCR, were also identified in the reviewed studies, mostly targeting vulnerable species like sharks and seahorses. Despite the great potential of HTS-based methods for authenticating seafood species, few studies have explored their utility in ESPO products. Notably, no research has yet explored the application of emergent molecular species identification technologies, such as CRISPR-based platforms, for seafood authentication in the ESPO. This relatively new technology has the potential to revolutionize seafood authentication, as indicated by recent studies reporting higher specificity and sensitivity levels, as well as shorter processing times compared to traditional DNA identification methods (Ying et al., 2022; Moorthy et al., 2025).

This study also conducted the first meta-analysis on the mislabeling of ESPO seafood products sold globally, revealing an overall mislabeling rate of 24.8%. As expected, the highest rates of mislabeling were detected in the final stages of the supply chain. A significant finding from this review is that threatened shark species were not only the most frequently mislabeled but also the most commonly encountered substitute species. This raises important concerns about conservation, fishery management, trade, and regulation of vulnerable resources. In light of this, laudable shark conservation initiatives, like the “Shark Free Ceviche” seal (Díaz-Ferguson et al., 2024), which has already been implemented in Colombia, should be replicated or adopted in more ESPO nations like Ecuador and Peru, which are major players in global shark fishing, trade, and consumption.

## Supplementary File legends

**Supplementary Table S1:** Metadata of samples included in the global mislabeling analysis of Eastern South Pacific seafood. Panel B shows detailed mislabeling rates results per each analyzed category

**Supplementary File S1:** R scripts used to determine confidence intervals

**Supplementary File S2:** R scripts used to analyze the statistical significance of mislabeling rates grouped by Continent

**Supplementary File S3:** R scripts used to analyze the statistical significance of mislabeling rates grouped by Country

**Supplementary File S4:** R scripts used to analyze the statistical significance of mislabeling rates grouped by Supply Chain Distribution Channel

**Supplementary File S5:** R scripts used to analyze the statistical significance of mislabeling rates grouped by Seafood Type

**Supplementary File S6:** descriptive analysis of the primers used in the reviewed articles

